# Effect of chlorination and pressure flushing of drippers fed by reclaimed wastewater on biofouling

**DOI:** 10.1101/2020.07.17.208074

**Authors:** Kévin Lequette, Nassim Ait-Mouheb, Nicolas Adam, Marine Muffat-Jeandet, Valérie Bru-Adan, Nathalie Wéry

## Abstract

Dripper clogging reduces the performance and service life of a drip irrigation system. The impact of chlorination (1.5 ppm of free chlorine during 1 h application) and pressure flushing (0.18 MPa) on the biofouling of non-pressure-compensating drippers fed by real reclaimed wastewater was studied at lab scale using Optical Coherence Tomography. The effect of these treatments on microbial composition (bacteria and eukaryotes) was also investigated by High-throughput DNA sequencing. Biofouling was mainly observed in inlet, outlet and return areas of the drippers. Chlorination limited biofilm development mainly in the mainstream of the milli-labyrinth channel. It was more efficient when combined with pressure flushing. Moreover, chlorination was more efficient in maintaining the water distribution uniformity. It reduced the bacterial concentration and the diversity of the dripper biofilms compared to the pressure flushing method. This method strongly modified the microbial communities, promoting chlorine-resistant bacteria such as *Comamonadaceae* or *Azospira*. Inversely, several bacterial groups were identified as sensitive to chlorination such as Chloroflexi and Planctomycetes. Nevertheless, one month after stopping the treatments the bacterial diversity re-increased and the chlorine-sensitive bacteria such as Chloroflexi phylum and the Saprospiraceae, Spirochaetaceae, Christensenellaceae and Hydrogenophilaceae families re-emerged with the growth of biofouling, highlighting the resilience of the bacteria from drippers. Based on PCoA analyses, the structure of the communities still clustered separately from never-chlorinated drippers, showing that the effect of chlorination was still present one month after stopping the treatment.

**Highlights:**

- The fouling of drippers is a bottleneck for drip irrigation using reclaimed wastewater
- Biofouling was lowest when chlorination was combined with pressure flushing
- The β-Proteobacteria and Firmicutes contain chlorine resistant bacteria
- The decrease of Chloroflexi by chlorination was transitory
- The bacterial community was resilient after the interruption of cleaning events

## 1. Introduction

Due to the scarcity of water resources, the use of reclaimed wastewater for crop irrigation is increasing significantly worldwide, particularly in arid and semi-arid countries (Ait-mouheb et al., 2020). Drip irrigation combined with the use of reclaimed wastewater has several advantages, as it provides water to the plant (e.g. roots) in optimal quantities and frequencies for plant growth (Goyal, 2018; Wang et al., 2013). The drippers are generally designed by a narrow labyrinth channel (internal flow section around 1mm) which makes the development of a turbulent regime possible composed of a main high velocity flow and vortex zones in channel corners (Al-Muhammad et al., 2019, 2016; Wei et al., 2012). This milli-labyrinth flow path can be clogged by suspended particles, chemical precipitation and biofilms (Green et al., 2018; Y. Li et al., 2019; Oliveira et al., 2017, 2020; Rizk et al., 2019; Zhou et al., 2018). The biofilm growth increases physical and chemical clogging (Li et al., 2013; Tarchitzky et al., 2013) and is considered a key factor in the clogging of drip irrigation systems using reclaimed wastewater (Song et al., 2017; Wang et al., 2017). Therefore, the evaluation of methods to control and limit the development of biofilms in these systems is necessary (Lamm et al., 2007).

Existing methods to reduce clogging include precipitation, filtration, chlorination, acidification and pressure flushing (Duran-Ros et al., 2009; Katz et al., 2014; Puig-Bargués et al., 2010; Song et al., 2017). Although chlorination has been recognised as the least expensive method to treat clogging due to biological growth, the chlorination schemes recommended in the scientific literature differ according to the type of reagent used (liquid sodium hypochlorite (NaOCl), calcium hypochlorite (Ca(OCl)_2_), gaseous chlorine (Cl_2_)) the concentration and injection interval application (Goyal, 2018; Rav-Acha et al., 1995) Indeed, chlorination can be applied from 0.4 ppm (Batista et al., 2009) to more than 100 ppm (Chauhdary et al., 2015). Both the chlorine concentration and the injection frequency had an impact on the effectiveness of reducing clogging. However the application of high chlorine concentrations might intensify the clogging by releasing clogging constituents that were previously stuck to the pipe wall (Rav-Acha et al., 1995) and can induce negative effects on crop growth, since a high chlorine concentration in the soil may lead to toxicity (Li and Li, 2009). Therefore, the use of low concentrations of free chlorine (1-2 ppm) at repeated frequencies (weekly, twice a week) has proven its effectiveness in limiting the fouling of the dripper (J. Li et al., 2010; Li et al., 2012; Song et al., 2017). Other studies have shown that the use of chlorine at too high frequencies (once or twice a week) can also promote the development of a biofilm resistant to chlorination (J. Li et al., 2010). However, the mechanisms and microorganisms involved in this process are little known. The in-depth analysis of the chlorination effects on the microbiome of drippers appears necessary to better understand the mechanisms of biofilm development and optimise the control of biofilm development (Zhou et al., 2017). Pressure flushing consists in washing water pipelines by increasing the hydraulic shear force within the pipe system. To be effective, flushing must be carried out often enough and at an appropriate rate to dislodge and transport accumulated sediment. The minimum flow velocity for flushing the drip irrigation system is 0.3-0.6 m s^−1^, thus removing clogging and particulate matter (Han et al., 2018; Lamm et al., 2007; Li et al., 2018). As for chlorination, the frequency of flushing (weekly to monthly) directly influences the efficiency of the method and the service life of the drip irrigation system (Lamm, 2013; Li et al., 2015, 2018; Puig-Bargués et al., 2010). Flushing the system as often as possible is recommended (Lamm et al., 2007), but this may promote the development of a flushing-resistant biofilm caused by specific bacteria (Li et al., 2015). Thus, despite studies aiming to optimise application parameters, the problem of clogging persists and the mechanisms involved are poorly understood.

The effectiveness of chlorination and pressure flushing methods on the biofouling of drip irrigation systems is often studied separately. However, several studies on drinking water systems have shown that combining the two methods was more effective in reducing the level of colonisation rather than using them separately (Mathieu et al., 2014; Tsvetanova, 2020). Thus, studying the effect of these methods alone and in combination will lead to a better understanding of the mechanisms involved.

The objectives of the present study were to determine the effect of chlorination and pressure flushing, combined or used alone on (1) biofilm kinetic development and (2) on the microbial communities in biofilms formed in irrigation systems fed by reclaimed wastewater (RWW). To evaluate if the effect of the treatments were transient or permanent, biofilms were also analysed 1 month after stopping the cleaning procedures. A non-destructive optical time-monitoring observation system was developed to study biofilm growth in commercial drippers (1 l h^−1^) by optical coherence tomography (OCT) (Lequette et al., 2020). The combined use of the OCT method and high-throughput sequencing made it possible to monitor the development of biofilm according to the cleaning methods used while evaluating the impact of these conditions on the bacterial composition. Although eukaryotes can influence the formation and development of biofilms (Parry, 2004; Parry et al., 2007), data on eukaryotes in drippers supplied by reclaimed wastewater are scarce (Dehghanisanij et al., 2005). Therefore, the impact of these cleaning methods on eukaryotes was also investigated.

## 2. Materials and Methods

### Experimental setup

#### Experimental setup and irrigation procedure

Non-pressure-compensating (NPC) drippers (model D2000, Rivulis Irrigation SAS, Lespinasse, France) with a flow rate of 1 l.h^−1^ (Table 1) were used. The NPC drippers were placed in a transparent tube (internal diameter 15 mm, TubClair, Vitry-le-François, France), allowing the analysis of the biofilm development along the channel over time without disturbances. More details are presented in Lequette et al. (2020). Four irrigation lines with nine dripper systems per line were placed on the test bench: control (C-without cleaning event), a pressure flushing (PF), chlorination line (Cl) and pressure flushing/chlorination (PFCl) (see Cleaning procedures for details). Each of the four lines was connected to a separate tank (total volume 60l) and a pump (JetInox 82M, DAB, Saint-Quentin-Fallavier, France) (Figure 1.B). A disk filter (mesh size 0.13mm) was installed to reduce the physical clogging of emitters following the technical recommendations for this type of dripper. The inlet pressure was set at 0.08 MPa (the dripper’s nominal working pressure) with a pressure gauge. A gutter, connected to each respective tank, was installed below each lateral line to collect the water discharged from the drippers during the experiments. The lines were supplied twice a day, five days out of seven for 1 h, with an interval of 6h off. Discharge measurements were performed each week to evaluate emitter performance and the clogging. The relative average discharge of drippers (Li et al., 2015) was used to assess the drip irrigation performance and was calculated in Equation 1:

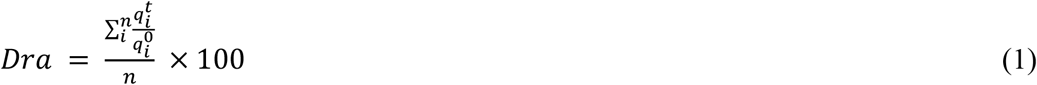

where q_i_^0^ indicated the nominal flow of drippers (l.h^−1^), q_i_^t^ the measured flow rate (l.h^−1^) and n was the total number of experimental emitters.

**Figure 1.**
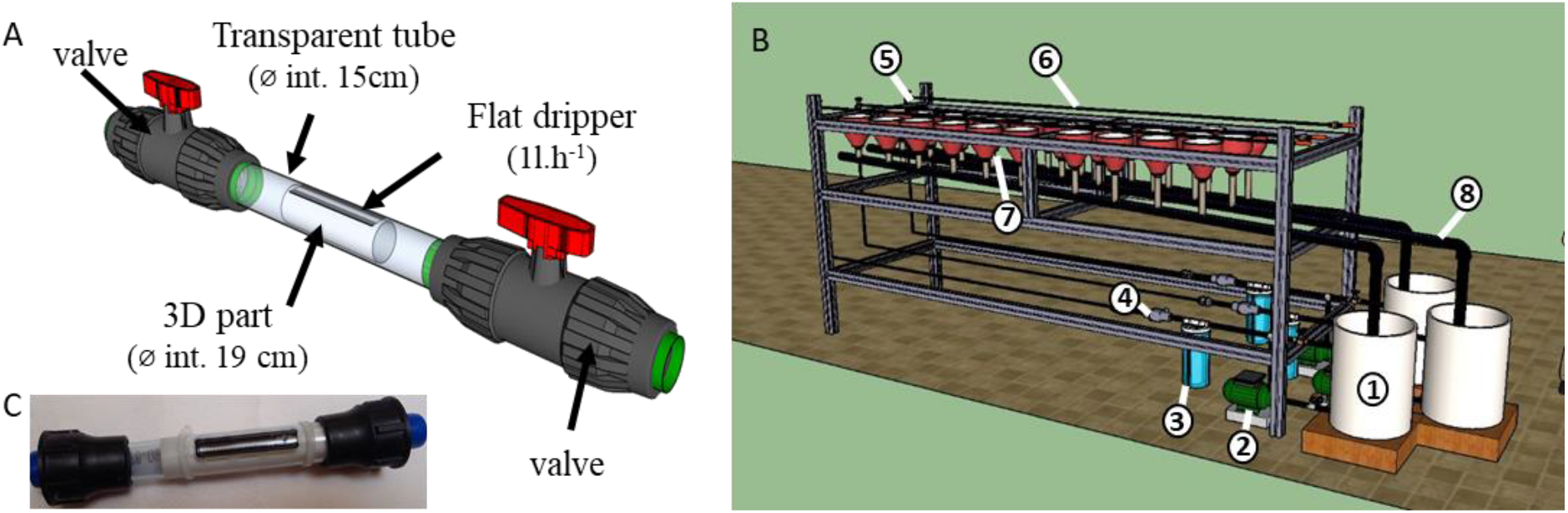
Dripper system (A, C) and test bench (B). The drippers were placed in a transparent tube to enable optical measurements. The test bench was composed of 1. a tank (60l); 2. a water pump; 3. a 0.13mm mesh screen filter; 4. a pressure reducer; 5. a pressure gauge; 6. the drip line with an emitter system located at 10 cm intervals; 7. a collector; 8. a gutter.

**Table 1.**
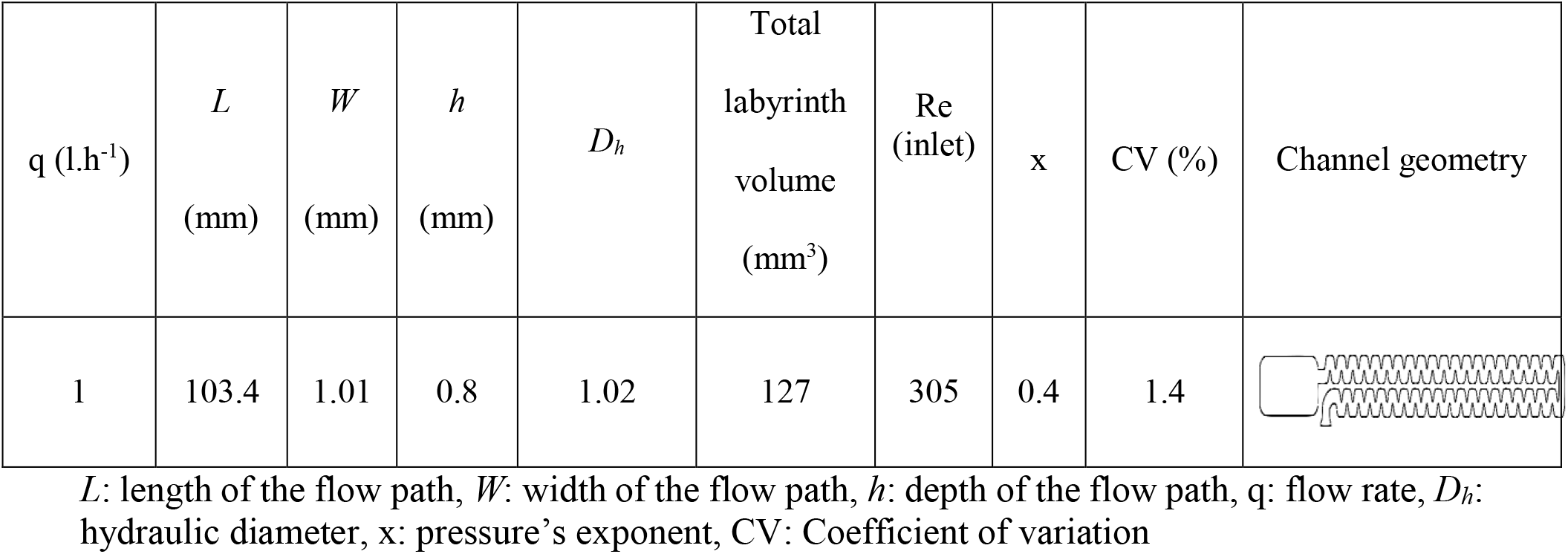
Dripper parameters.

| q ($\text{l.h}^{-1}$) | L<br>(mm) | W<br>(mm) | h<br>(mm) | $D_h$ | Total<br>labyrinth<br>volume<br>($\text{mm}^3$) | Re<br>(inlet) | x | CV (%) | Channel geometry |
| --- | --- | --- | --- | --- | --- | --- | --- | --- | --- |
| 1 | 103.4 | 1.01 | 0.8 | 1.02 | 127 | 305 | 0.4 | 1.4 |  |

L: length of the flow path, W: width of the flow path, h: depth of the flow path, q: flow rate, $D_h$ : hydraulic diameter, x: pressure's exponent, CV: Coefficient of variation

The drippers were considered clogged when the outflow presented a discharge of less than 75% (Yu et al., 2018b).

Table 1 lists the dripper characteristics. The flow regimes were characterised by a Reynolds number (Re). The Reynolds number (Re) was computed using the following Equation 2:

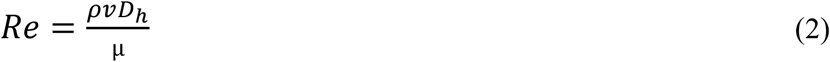

where *ρ* is water density (kg.m^−3^), *v* the water velocity across the pipe (m.s^−1^), *D*_*h*_ the hydraulic diameter (m), and μ the water dynamic viscosity (Pa s).

The hydraulic diameter), *D*_*h*_, was calculated for a rectangular cross-section as (Equation 3):

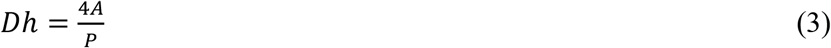

where A: area (mm) and P for the perimeter (mm).

#### Physicochemical and microbiological quality of TWW

The irrigation lines were supplied with reclaimed wastewater from the Murviel-Les-Montpellier treatment plant in the South of France (43.605034° N, 3.757292° E). The wastewater treatment plant is designed around stabilisation ponds with three successive lagoons (13 680 m^3^, 4784 m^3^ and 2700 m^3^) and a nominal capacity of 1,500 Inhabitant Equivalent. RWW was placed in a 60l tank and changed twice a week to maintain the quality close to that of the wastewater from the treatment plant. Each week (n=16), several physical-chemical and microbiological analyses were performed to evaluate the RWW quality. Chemical oxygen demand (COD), ammonia, nitrate, and phosphorus concentrations (mg l^−1^) were measured with a spectrophotometer (DR1900, Hach Company, Loveland, CO, USA) using LCK Hach reagents^®^. Conductivity and pH were measured with probes (TetraCon^®^ 925 and pH-Electrode Sentix^®^ 940, WTW, Wilhelm, Germany). Faecal coliforms, *E. coli,* and *Enterococci* were quantified using the IDEXX method (Colilert18 and Enterolert, IDEXX Laboratories, Westbrook, ME) according to the supplier’s recommendations. The main effluent properties are listed in Table 2.

**Table 2.** Physico-chemical and microbiological characteristics of the RWW.

| Characteristics | Units | Mean<br>(n=9) | SD |
| --- | --- | --- | --- |
| COD (LCK 614) | $\text{mg l}^{-1}$ | 159.7 | 25.1 |
| Total suspended solids | $\text{mg l}^{-1}$ | 65.9 | 13.8 |
| Ammonia (LCK 303) | $\text{mg l}^{-1}$ | 28.1 | 11.8 |
| Nitrate (LCK 339) | $\text{mg l}^{-1}$ | 3.1 | 1.1 |
| Phosphorus (LCK 350) | $\text{mg l}^{-1}$ | 7.0 | 0.8 |
| Conductivity | $\mu\text{S cm}^{-1}$ | 1319.3 | 82.1 |
| Dissolved oxygen | $\text{mg l}^{-1}$ | 8.8 | 0.4 |
| pH |  | 7.9 | 0.1 |
| Total coliforms | MPN/100<br>mL | 3.10E+05 | 1.60E+05 |
| *Escherichia coli* | MPN/100<br>mL | 1.08E+05 | 0.8E+05 |
| *Enterococci* | MPN/100<br>mL | 2.73E+04 | 1.72E+04 |

### Cleaning procedures

Cleaning event tests were divided in two periods: a cleaning period (71 days) followed by a period without cleaning events (33 days) in order to study the interruption of these treatments on the biofilm’s regrowth. During the cleaning period, chlorination and pressure flushing were applied once a week during 1h.

For the disinfection method, liquid sodium hypochlorite (NaOCl) with a residual chlorine concentration >9% was used for chlorination. NaOCl was added directly to the tank in order to reach 1.5 ppm of free chlorine at the end of the drip irrigation line. After the system had been running for 1h, the residual free chlorine concentration was tested every 10 min at the end of the drip irrigation lines using LCK 310 Hach reagents^®^. Depending on the measurements, NaOCl was added in the tanks to maintain the 1.5 ppm of free chlorine in the system.

The maximum working pressure used for the drippers (1 l.h^−1^) was 0.25MPa according to the manufacturer’s recommendations. However, the use of the transparent tubes around the dripper systems did not allow this pressure to be reached without damaging the system. Therefore, for lateral flushing procedures, the valves at the end of the drip-line were closed and the pressure was increased to 0.18MPa. The pressure was controlled using a manometer placed at the inlet of the dripline. After each cleaning event, RWW from the test bench was removed and replaced by new RWW.

### Image acquisition and processing

#### Image acquisition

Optical coherence tomography (OCT) was used to study the kinetics of biofilm in the drippers and along the milli-channel throughout the experimental period (Table 3) after 3 weeks of running. Measurements were made in situ and non-invasively through the transparent tube. For the measurements, the valves of the dripper systems were closed to keep the drippers in the water. The dripper system was then disconnected from the irrigation line. Measurements were taken at least once every two weeks. Due to the number of drippers, the monitoring of the 36 drippers (9 per lines) by OCT was carried out over one week. After the OCT measurement, each dripper was returned to its original location on the test bench. The three-dimensional OCT measurements were acquired using a Thorlabs GANYMEDE II OCT (LSM03 lens, axial resolution= 2.1 μm, lateral resolution= 8μm; Thorlabs GmbH, Lübeck, Germany). The size of the axial voxel in water (n = 1.333) of the GANYMEDE II is 2.1 μm. OCTs have a center wavelength of 930 nm.

**Table 3.**
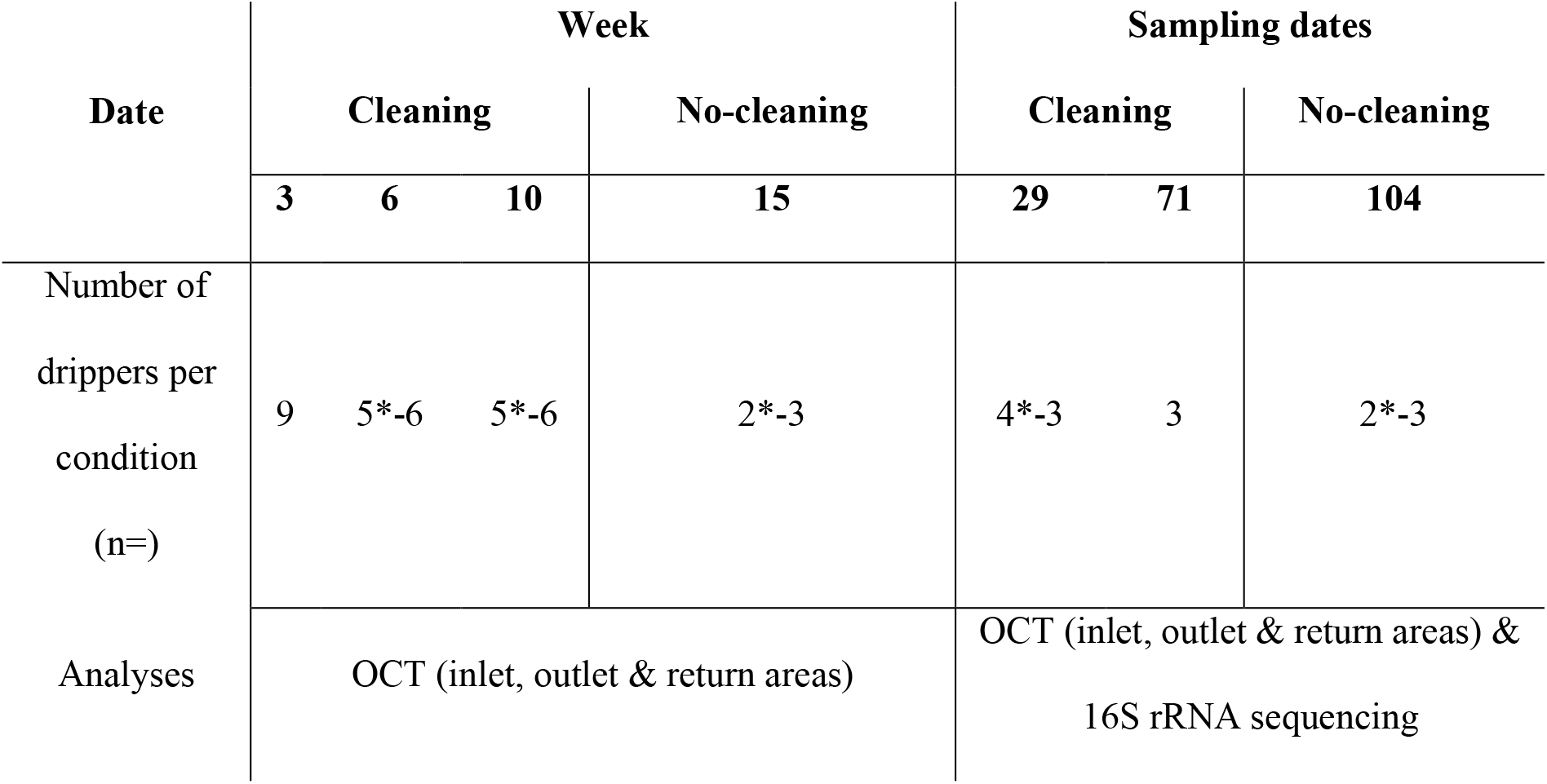
Analysis and sampling schedule. *: After one month, one dripper from the control line was totally clogged. After 29 days this dripper was sampled along with 3 randomly chosen ones. This explains why the number of drippers monitored and sampled at 104 days is lower than in the other conditions.

| Date | Week |  |  |  | Sampling dates |  |  |
| --- | --- | --- | --- | --- | --- | --- | --- |
|  | Cleaning |  |  | No-cleaning | Cleaning |  | No-cleaning |
|  | 3 | 6 | 10 | 15 | 29 | 71 | 104 |
| Number of drippers per condition (n=) | 9 | 5*-6 | 5*-6 | 2*-3 | 4*-3 | 3 | 2*-3 |
| Analyses | OCT (inlet, outlet & return areas) |  |  |  | OCT (inlet, outlet & return areas) & 16S rRNA sequencing |  |  |

#### Image analysis

OCT acquisition was performed in order to follow the biofouling development in time (volume and thickness) depending on the treatment used. First, 3-D OCT datasets were processed in Fiji (running on ImageJ version 1.50b, Schindelin et al. (2009)) and converted into 8-bit grayscale. The datasets were resliced from top to bottom into image stacks and regions of interest (inlet, outlet and return areas) were selected (Figure S1 in Supplemental material). The remaining parts were allocated to the background (black). Secondly, an in-house code was used to detect the pixels associated with the plastic tube and removed using MATLAB R2018r (MathWorks ®, version 2018b). A threshold (adapted to each dataset) was applied to binarize the dataset and the region above the interface was quantified as biofilm. For each position (x, y), the pixels associated with the biofilm (up to the threshold) were summed (on z) to obtain the thickness of the biofilm. The biofilm volume (mm^3^) was calculated for each area according to Equation 4:

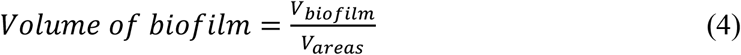

where *V*_*biofilm*_ is the biofilm volume and *V*_*areas*_ is the volume of the area of interest (inlet, outlet, return).

After the first visualisation of the data, an additional data-driven approach was realised conducting to apply the quantification of the biofilm along the labyrinth divided into two zones (around the edges / around the middle). Zones were created using a ‘median split’ procedure. The width of the tube was determined for each x-step and divided by four in order to separate the tube into two equivalent areas: i) the middle zone corresponds to the two central quarters, ii) the edge zone corresponds to the two side quarters summed.

### Analysis of microbial communities

#### Biofilm and RWW samplings

After the first visualisation of the data, an additional data-driven approach was realised conducting to apply the quantification of the biofilm along the labyrinth divided into two zones (around the edges / around the middle). Zones were created using a ‘median split’ procedure. The width of the tube was determined for each x-step and divided by four in order to separate the tube into two equivalent areas: i) the middle zone corresponds to the two central quarters, ii) the edge zone corresponds to the two side quarters summed.

#### DNA extraction

DNA was extracted using the PowerWater® DNA Isolation Kit (Qiagen, Hilden, Germany). The samples (drippers or filters) were placed in 5 ml tubes containing beads. The manufacturer’s instructions were then followed. The DNA concentration was measured and the purity checked by spectrophotometry (Infinite NanoQuant M200, Tecan, Austria). The extracted DNA was stored at −20°C.

#### Bacterial quantification by qPCR

Total bacterial quantification was performed by qPCR on biofilms from drippers targeting the V9 region from 16S rDNA. The amplification reactions were performed in triplicate, and at two dilutions to check for the absence of inhibition of the PCR reaction. Reaction mixes (12μl) contained 2.5μl of water, 6.5μl of Super Mix qPCR (Invitrogen), 100nM forward primer BAC338 (5’-ACTCCTACGGGAGGCAG-3’), 250nM of reverse primer BAC805 (5’-GACTACCAGGGTATCTAAT CC-3’) and 50nM of probe BAC516 (Yakima Yellow-TGCCA GCAGC CGCGG TAATA C –TAMRA) (Yu et al., 2005). The cycling parameters were 2 min at 95°C for pre-incubation of the DNA template, followed by 40 cycles at 95°C for 15 sec for denaturation and 60°C for 60 sec for annealing and amplification.

#### Illumina sequencing

The PCR amplified the V4-V5 region of 16S rRNA genes with 30 cycles (annealing temperature 65°C) using the primers 515U (5′-GTGYCAGCMGCCGCGGTA-3′) and 928U (5′-CCCCGYCAATTCMTTTRAGT-3′) (Wang and Qian, 2009). A PCR amplified the 18S rRNA genes (30 cycles, annealing temperature 56°C) was also performed using the primers 5’-GCGGTAATTCCAGCTCCAA-3’ and 5’-TTGGCAAATGCTTTCGC-3’ (Hadziavdic et al., 2014) on dripper biofilms at the end of the cleaning period. Adapters were added for multiplexing samples during the second amplification step of the sequencing. The resulting products were purified and loaded onto the Illumina MiSeq cartridge for sequencing of paired 300 bp reads according to the manufacturer’s instructions (v3 chemistry). Sequencing and library preparation was performed at the Genotoul Lifescience Network Genome and Transcriptome Core Facility in Toulouse, France (get.genotoul.fr). Mothur (version 1.39.5) (Schloss et al., 2009) was used to associate forward and reverse sequences and clustering at four different nucleotides over the length of the amplicon. Uchime (Edgar et al., 2011) was used to identify and remove chimera. Sequences that appeared less than three times in the entire data set were removed. In all, 16S rRNA sequences were aligned using SILVA SSURef NR99 version 128 (Schloss et al., 2009). Finally, sequences with 97% similarity were sorted into operational taxonomic units (OTUs) (Nguyen et al., 2016). The chloroplast sequences from 16S rRNA sequences were removed from the raw data and represented respectively 2.8% and 16.6% of the sequences in biofilms and in RWW samples. Finally, BLAST (http://www.ncbi.nlm.nih.gov/BLAST/) was used to locate publicly available sequences closely related to the sequences obtained from the samples. A total of 2,281,644 reads were grouped in 3210 OTUs at the 97% similarity level. The rarefaction curves indicated that the sequencing depths of all samples were adequate (Figure S2 in Supplementary material).

### Statistical analyses

Non-parametric statistical tests (Kruskal and Wilcoxon tests) were performed to compare the biofilm volume and the biofilm distribution along the labyrinth depending on the treatment used. The Shannon diversity index, the reciprocal Simpson index, and the nonparametric richness estimators Chao1 was also calculated for each RWW and biofilm sample. Chao1 richness estimates were based on singletons and doubletons as described by Chao1 (Chao, 1984). Kruskal-Wallis tests were performed to compare indices of diversity and richness according to the origin of the sample, and biofouling rate between the drippers. For the comparison of bacterial community structures, a dissimilarity matrix (Bray-Curtis) was performed and visualised using principal coordinate analysis (PCoA). A one-way analysis of similarities (ANOSIM) was used to identify significant differences in community assemblage structure between samples based on the origin of the sample (Clarke, 1993). The genera that contribute most to the divergence between two habitats were identified using Similarity Percentage (SIMPER) analysis (Clarke, 1993). The sequencing data analysis was processed under R v3.4 (www.r-project.org) through R-Studio (http://www.rstudio.com/) with the phyloseq package (McMurdie and Holmes, 2012).

## 3. Results

The development kinetics of the biofilms under chlorination and pressure flushing treatments, combined or used alone, were followed for 71 days using OCT measurements and compared to each other and to non-treated controls. The drippers were collected after 29 and 71 days in order to characterise the effect of these treatments on the bacterial composition of the biofilms. After 71 days, the treatments were stopped and drippers were analysed at day 104 to evaluate if the effect of the treatments were permanent or transitory.

### Biofouling kinetic analysis

#### Dynamic changes for outflow ratio variation of drippers

Changes of discharge ratio (Dra) along the lateral are shown in Figure 2. Dra decreased mainly for control (C) and pressure flushing (PF) lines over time, reaching about 80% after 2 months. The Dra of the chlorinated lines (Cl and PFCl) was close to 100% after 2 months of treatment, meaning that chlorination allowed a better control of the flow rates than purge only. For the same treatment, the Dra sometimes increased between two measurements, which indicates that the clogging of the drippers can be temporary and that detachment can occur. In the control line, the majority of the drippers (7/9 after 71 days) presented a discharge of less than 75% and one dripper was totally clogged after one month (no outflow). In the pressure line, 4/9 drippers were considered as clogged after 71 days while none were clogged for chlorinated lines (chlorination alone or combined with pressure flushing). After 1 month without any cleaning steps, the Dra from the C and PF lines increased but were still lower than the Dra from the chlorinated lines.

**Figure 2.**
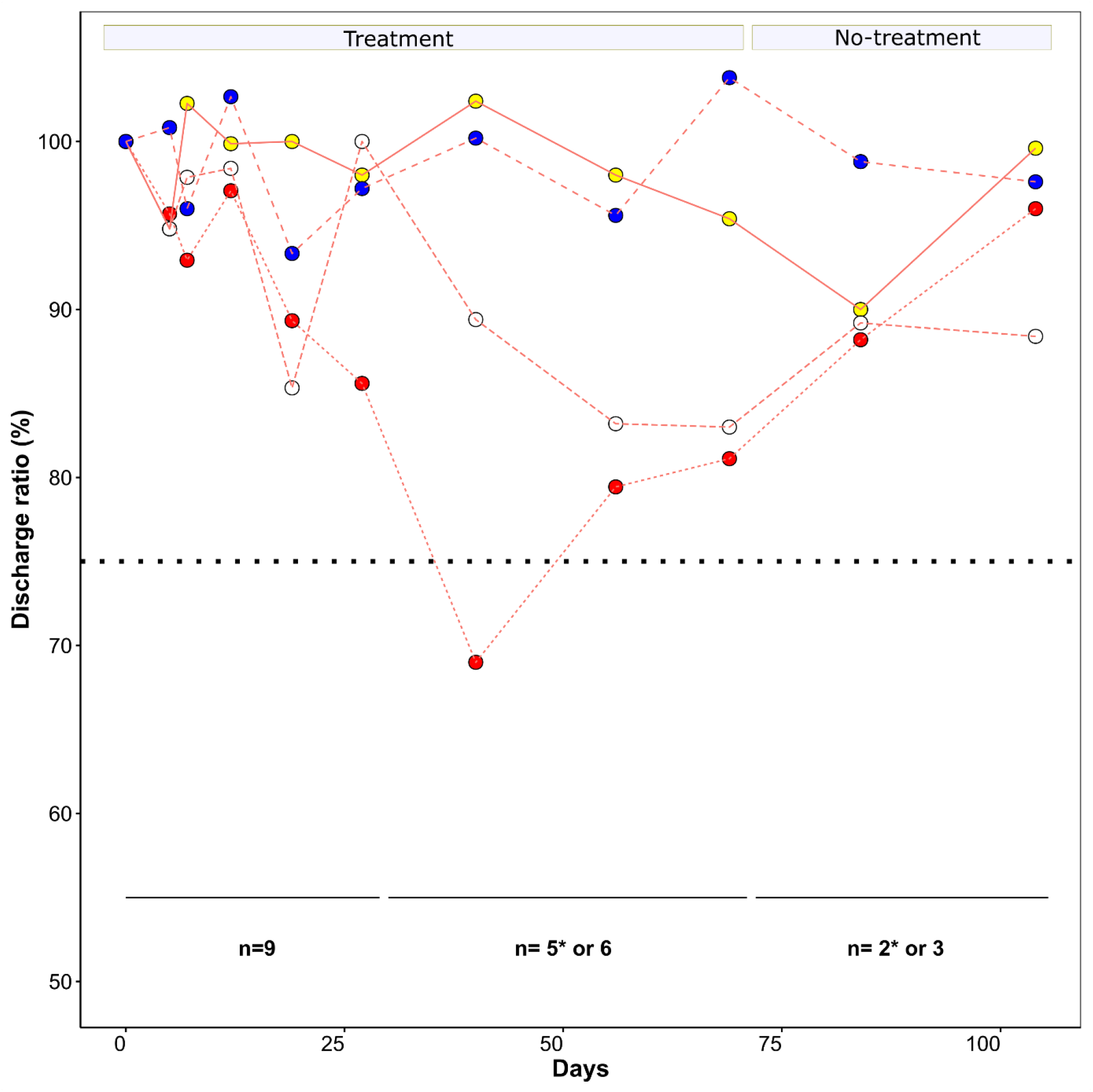
Discharge ratio variation according to the cleaning procedure used. Control 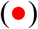, Pressure Flushing (○), Chlorination 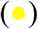 and Pressure Flushing combined with Chlorination 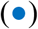. The drippers were considered as clogged when the discharge was less than 75% (dotted line). *n* corresponds to the number of drippers for each line (*: data control line).

#### Chlorination combined with pressure flushing decreased biofouling

The volume and thickness of the biofilm were determined using OCT in the areas of the canal most sensitive to fouling: the inlet, outlet and return zones.

The biofilm volume increased during the cleaning phase (p-value < 0.05, Figure 3). Although the average volume was lower in the PFCl condition after 10 weeks, there were no statistically significant differences between conditions indicating that the treatments had no significant effect on the overall biofouling level. This may be due to important variability between the triplicates. After one month without cleaning steps (at week 15), the mean biofouling volume increased in PF, Cl and PFCl conditions and volume was higher in the PF condition than in PFCl (p-value = 0.06) (Figure 3).

**Figure 3.**
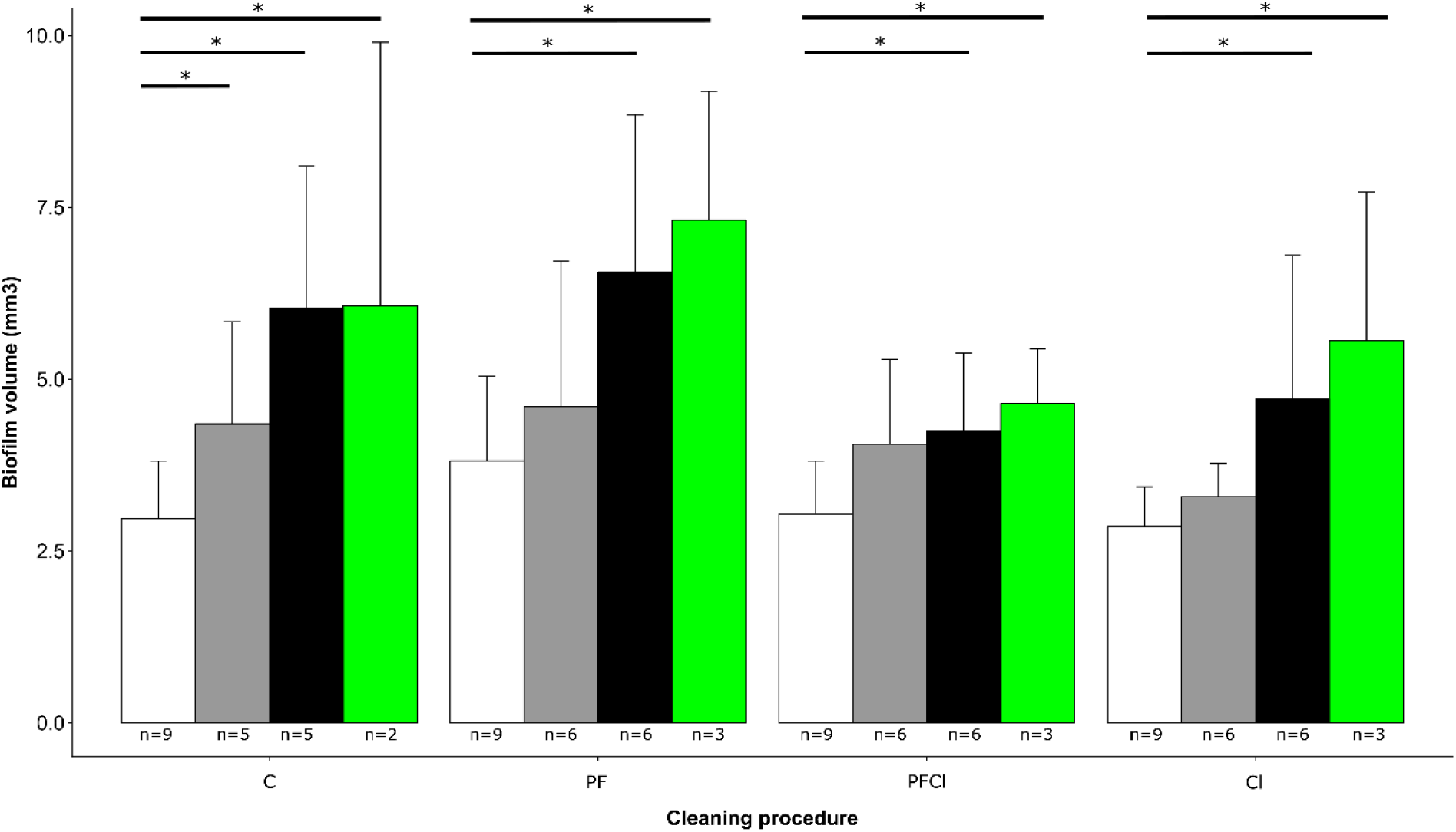
Evolution over time of the biofilm volume according to the cleaning method used. (at weeks 3, 6, 10 (cleaning phase) and 15 (no-cleaning phase); white, grey, dark grey and green respectively). C= control, PF= Pressure flushing, PFCl= Pressure flushing combined with chlorination, Cl= Chlorination. *n* refers to the number of drippers. The biofouling volume was quantified as the sum of the fouling volumes developed for the inlet, return and outlet areas. * shows significant differences (p-value < 0.05) on the conover test.

The study of the volume of clogging in the areas of interest shown that the biofouling tended to be higher in the inlet areas whatever the treatment was (Figure S3). Moreover, the outflow of the dripper was negatively correlated to the increase of the inlet biofouling volume after 10 weeks (Pearson’s test, r=-0.44, p<0.05), which may explain the decrease in Dra observed for lines C and PF (Figure 2).

Although the overall volume of biofilm in the drippers and sensitive areas was statistically similar over time between the conditions, the thickness of the biofilm was significantly influenced by the treatment used. Figure 3 shows the increase in biofouling thickness in the inlet areas during the cleaning period at weeks 3, 10 and 15. The inlet of the channel was the most sensitive area, in particular in the first baffle (p-value <0.05) where the average thickness increased by more than 0.5 mm (depth of the channel: 0.8 mm). The biofouling of the C drippers increased along the labyrinth but did not exceed 0.3-0.4 mm (Figure 4). OCT images showed that biofouling thickness decreased mainly in the middle of the channel when chlorination was applied (Cl and PFCl lines) at the inlet (Figure 4), outlet (Figure S4) and close to the return area (Figure S5) compared to C and PF lines.

**Figure 4.**
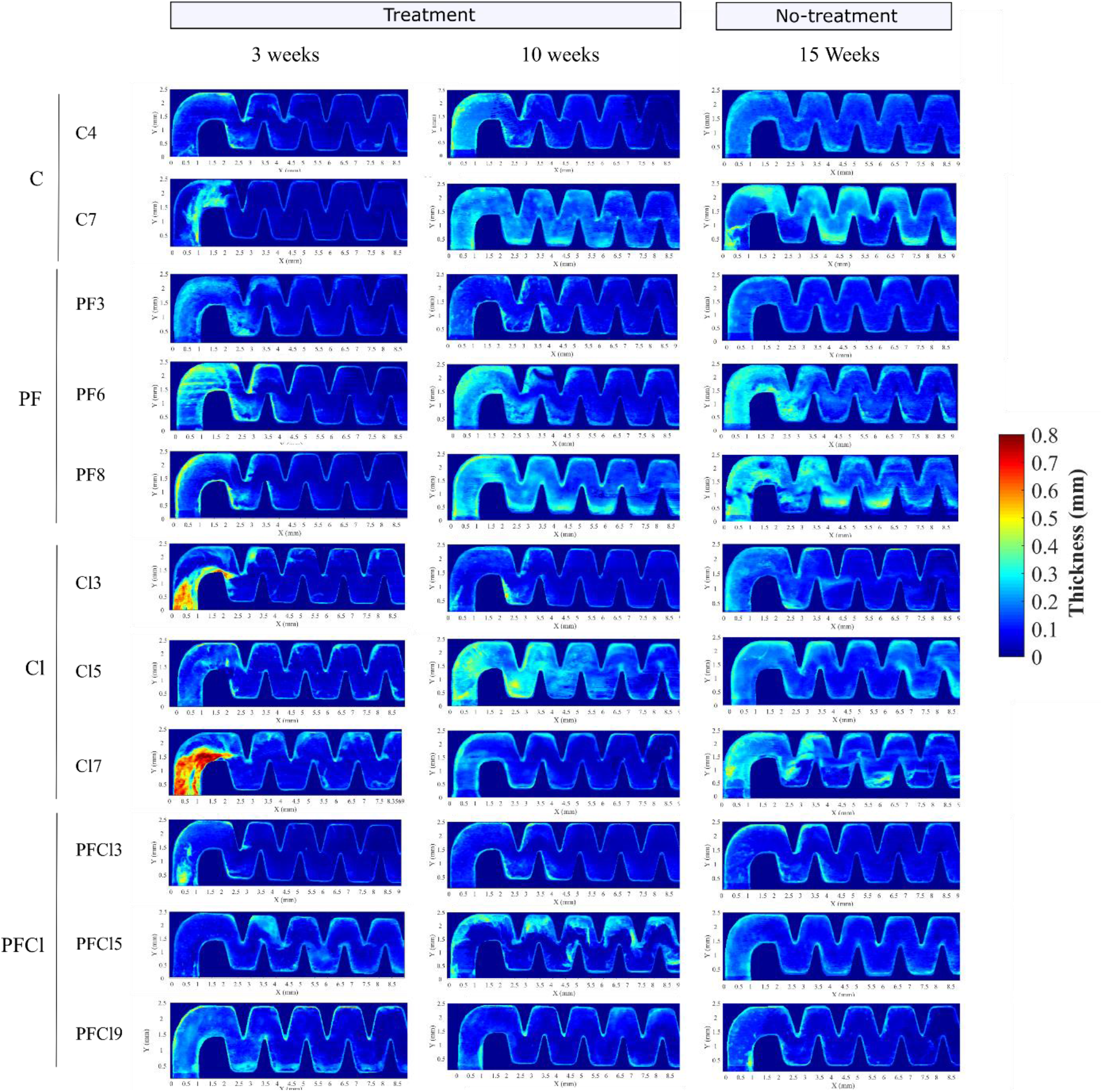
Biofilm thickness at the inlet of drippers under C (Control), PF (Pressure flushing), Cl (Chlorination) and PFCl (Pressure flushing/Chlorination) conditions measured after 3, 10 and 15 weeks. The drippers presented are those also analysed by high-throughput sequencing at 104 days.

To further explore the differences in locations of the biofilm between conditions, the inlet area was divided along the labyrinth into 2 parts: the middle and the edges. The number of pixels associated with biofouling was then determined in each of these areas. Figure 5 shows the mean pixels quantity of biofouling along the inlet channel in the middle and along the edges at the end of the cleaning step (10 weeks). When chlorination was combined with pressure flushing (PFCl), the number of pixels associated with biofouling along the inlet channel was statistically lower in the middle and close to the edges compared to the C, PF and Cl lines. Biofouling tended to be lower in the middle part than in the edges part. After 1 month without cleaning events, the number of pixels associated with biofouling along the inlet channel was similar in the middle and on the edges between all tested conditions (Figure S6). This means that the use of chlorination combined with increasing pressure, affected the location of the biofilm (edges or middle zone) rather than the overall amount of biofouling in the inlet zone. Moreover, this means that the combination of chlorination and pressure flushing was more efficient than the chlorination alone to limit biofilm growth.

**Figure 5.**
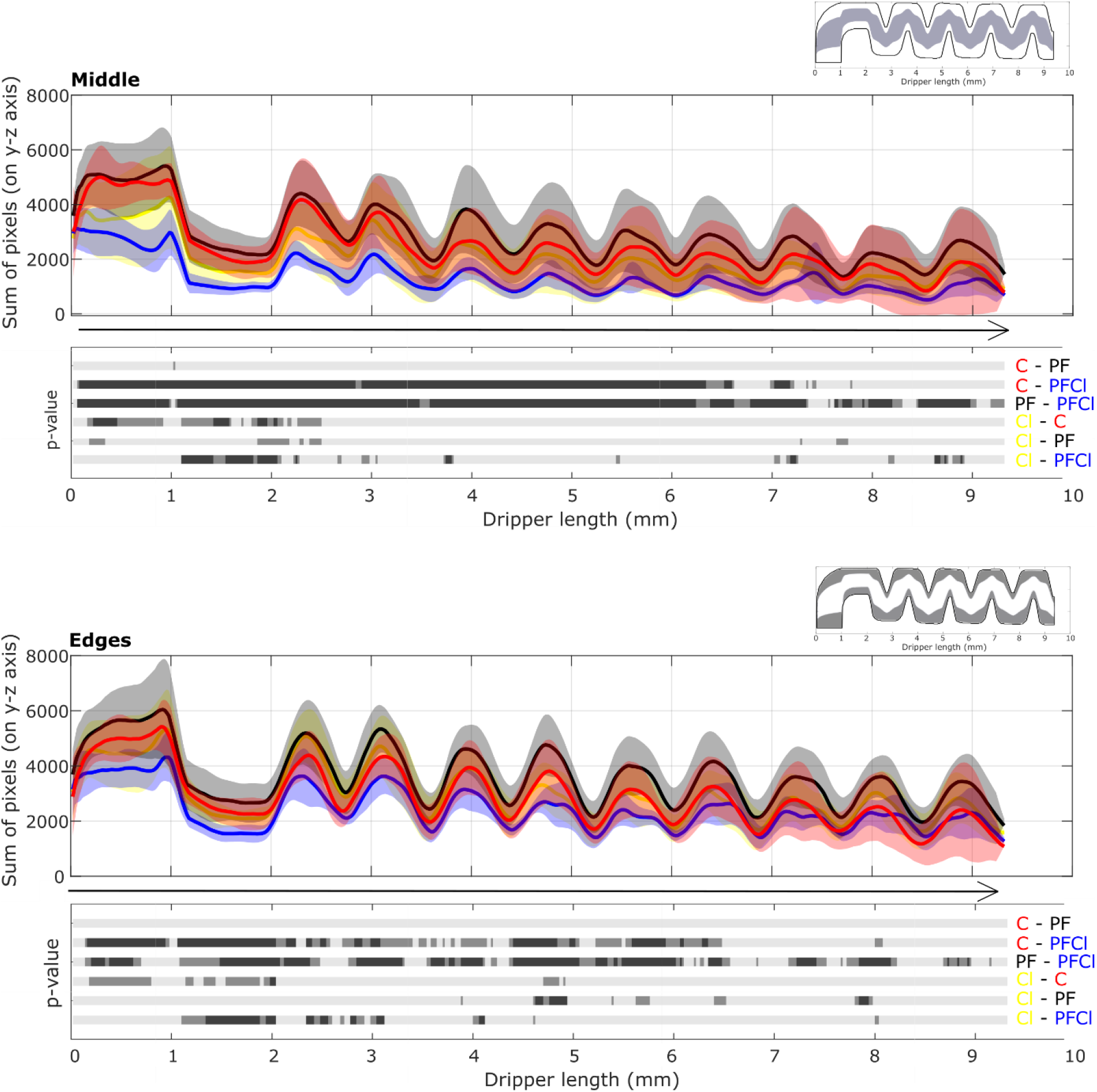
Means and standard deviation of pixels associated with biofouling in the middle and on the edges of the inlet dripper channel after 10 weeks of treatment. For each position of x, pixels in y-z are summed. Grey areas on inlet schemes at right-top indicate the zone of interest. Control (C-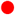), Pressure flushing (PF-●), Chlorination (Cl-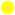) and Pressure flushing combined with Chlorination (PFCl-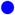); the arrow indicates the direction of the flow along the channel; n=6 per condition. P-value graphs show the results of the Wilcoxon tests with 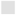: non-significant, 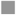: p<0.1, ■: p<0.05.

For some chlorinated drippers (Cl3, Cl7), the thickness between the 19^th^ and 71^st^ days had decreased around the first inlet baffle, suggesting that some detachment of biofilm occurred (Figure 4).

#### Biofouling level and bacterial quantification

The quantity of bacterial cells in the drippers during the cleaning period was between 10^9^ and 10^10^ copies of 16S rDNA at 29 days and between 10^10^ and 10^11^ copies at 71 days (Figure 6A) (background level of 10^3^ copies of 16S rDNA at t0). The number of copies of 16S rDNA increased significantly between days 29 and 71 for the C, PF and PFCl conditions (p-value<0.05) but not for the Cl condition (p-value>0.05). At 29 days, the number of copies of 16S rDNA in the control (C) and Cl conditions were higher than in the PFCl condition (p-value <0.05). At 71 days, the number of copies of 16S rDNA remained higher in the C condition than in the PFCl condition, which means that the combination of chlorination and pressure flushing reduces the number of bacteria more efficiently than pressure flushing (PF) or chlorination alone (Cl). Regardless of the protocol studied, the fouling level given by the volume of biofilm was positively correlated to the number of bacteria during the cleaning period (Pearson test, r=0.66) at 71 days, meaning that the number of bacteria in the dripper increased with the biofouling (Figure 6B).

**Figure 6.**
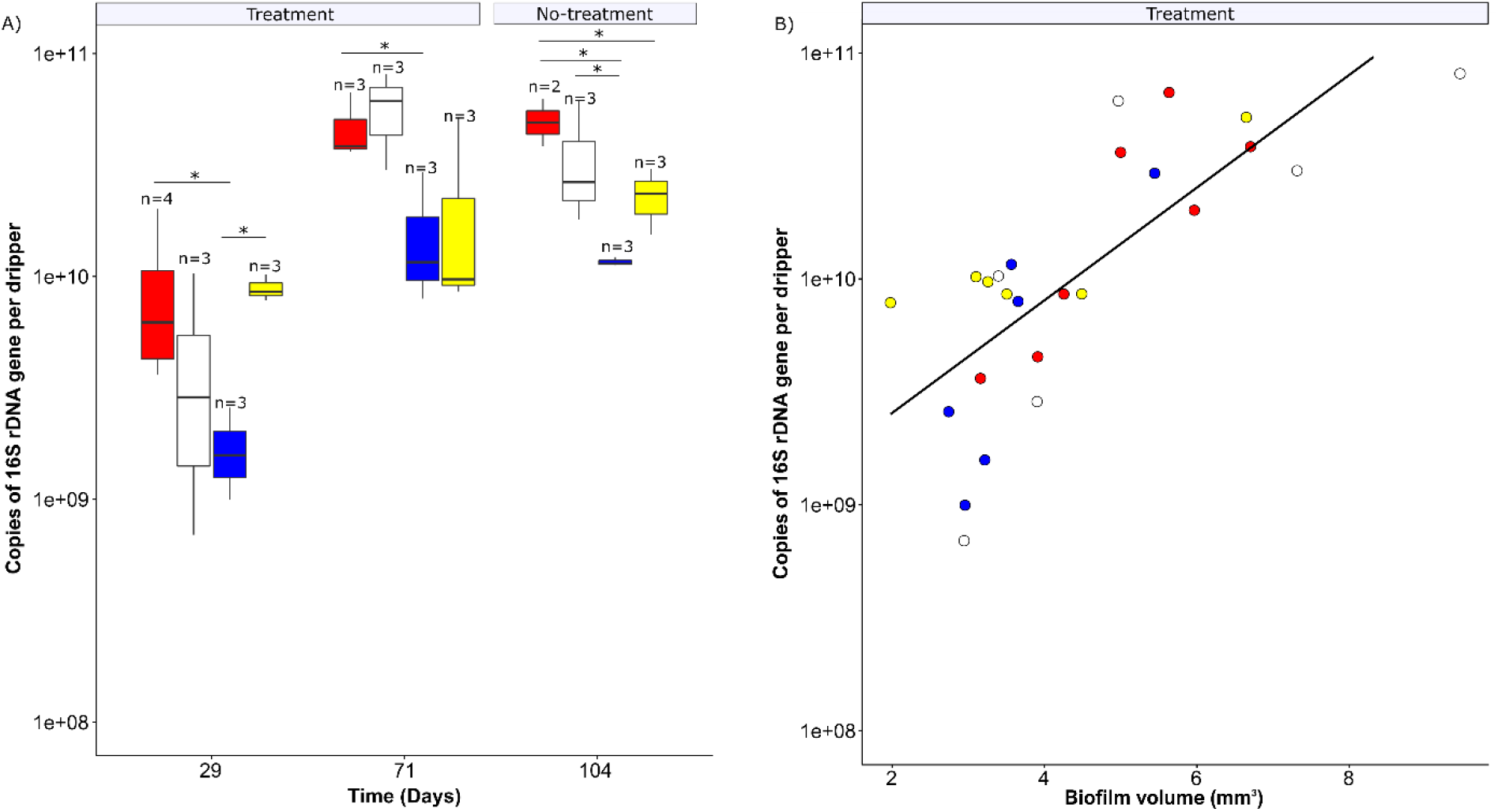
Evolution of the bacterial quantity by dripper according to the cleaning method used (left) and the biofilm volume (right). Control 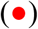, Pressure flushing (○), Chlorination 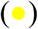 and Pressure flushing combined with Chlorination 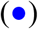. The biofouling volume was quantified as the sum of the fouling at the inlet, return and outlet areas. * indicates significant differences (conover-test, p-value < 0.05).

After one month without cleaning events, the number of copies of 16S rDNA was similar with that of the end of the cleaning period for each condition and the concentrations remained higher in the C and PF conditions compared with the PFCl condition.

### Bacterial Community Structure Analysis

The bacterial communities in dripper biofilms collected after 29 and 71 days were compared using 16S rDNA Illumina sequencing to evaluate the treatment effects. They were also compared with the communities present in reclaimed wastewater used to supply the dripper systems and renewed every week. After 71 days, treatments were stopped and biofilms were sampled again at 104 days to investigate the resilience of the communities, i.e. to test if the biofilms would then become closer to the control condition.

#### Structure of bacterial communities

During the cleaning period, the richness increased significantly between 29 days and 71 days for all conditions (Table 4, conover test, p-value<0.05) along with the biofouling level. The diversity indices remained similar (conover test, p-value>0.05) except for the Cl condition where the diversity indices increased after 71 days.

**Table 4.**
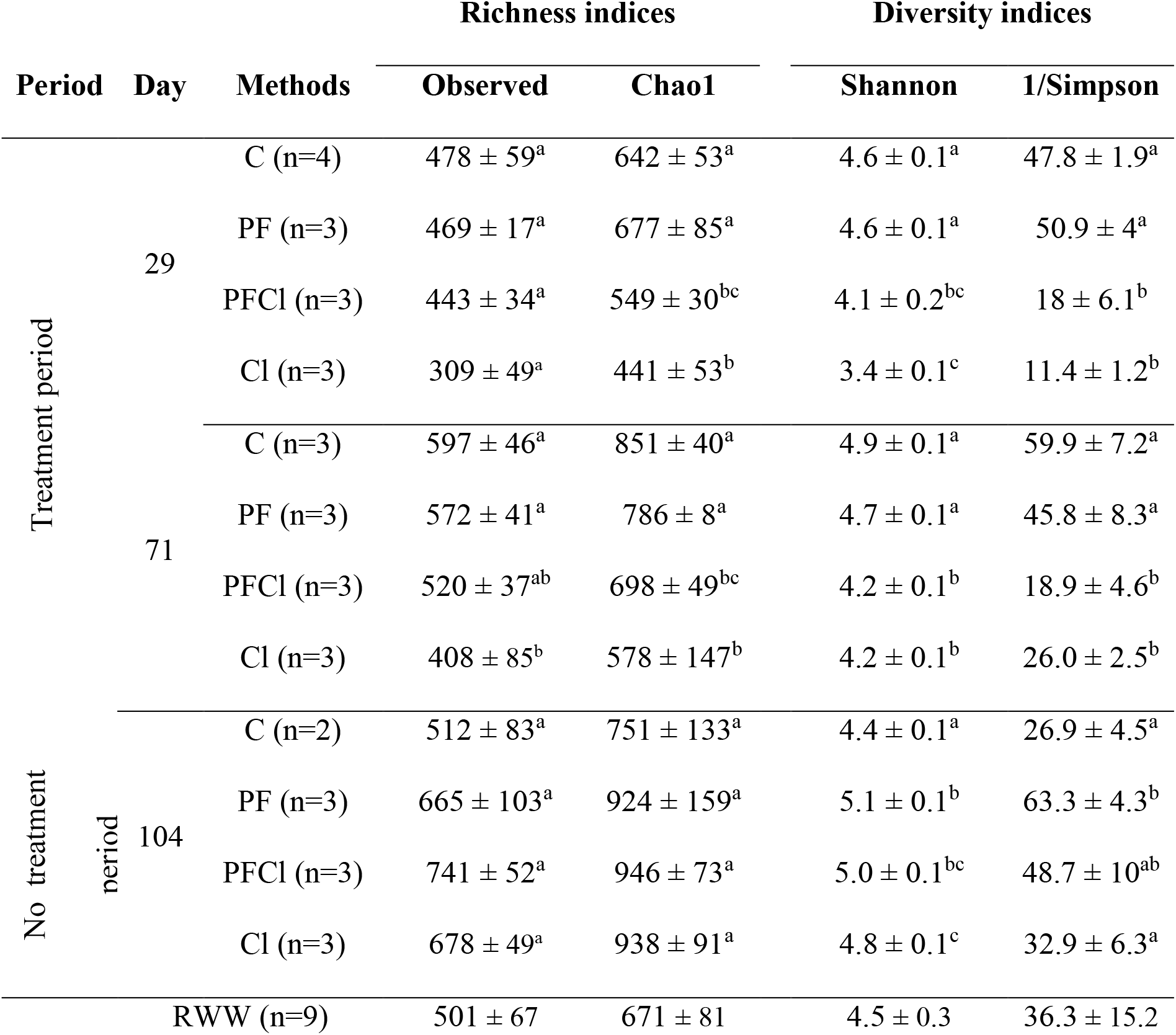
Richness and diversity indices according to the cleaning method.

The richness and diversity indices from the C and PF conditions were higher than in both chlorinated conditions (Cl and PFCl) at 29 days and 71 days (Table 4) indicating that the structure of the bacterial community was impacted by the treatment used. After the cleaning period (104 days), the richness and diversity indices were similar between the drippers of the different lines. However, the richness had increased in the Cl and PFCl lines while it remained stable in the C and PF lines. The Shannon index increased in the PF, Cl and PFCl conditions and was statistically higher than in the control (C) condition. In addition, the 1/Simpson’s index for the C line decreased from 71 days to 104 days.

Kruskal test and the conover ad hoc test were performed for each sampling time; the letters show the results of the conover test.

PCoA confirmed that the bacterial community changed depending on the cleaning method and over time (Figure 7). Both the cleaning method and time were significant factors for explaining the differences in community structure (p<0.05, Adonis). Pressure flushing, compared to the control condition, had no effect on the bacterial community structure. This was also true for pressure flushing combined with chlorination (when compared to chlorination alone). Thus, during the cleaning period, the bacterial population clustered depending on the use of chlorine (Adonis, p<0.05, R^2^_29 days_=0.58 and R^2^_71 days_= 0.68). At 29 days, the bacterial populations from the C dripper biofilms were significantly different from the Cl and PFCl dripper biofilms (Adonis, p<0.05, R^2^_C-Cl_=0.73 and R^2^_C-PFCl_=0.72 respectively). After 1 month without cleaning events, the clustering depending on the use of chlorine was maintained (Adonis, p<0.05, R^2^_104 days_=0.46) but there was no statistical difference between the chlorinated and non-chlorinated conditions (Adonis, p>0.05).

**Figure 7.**
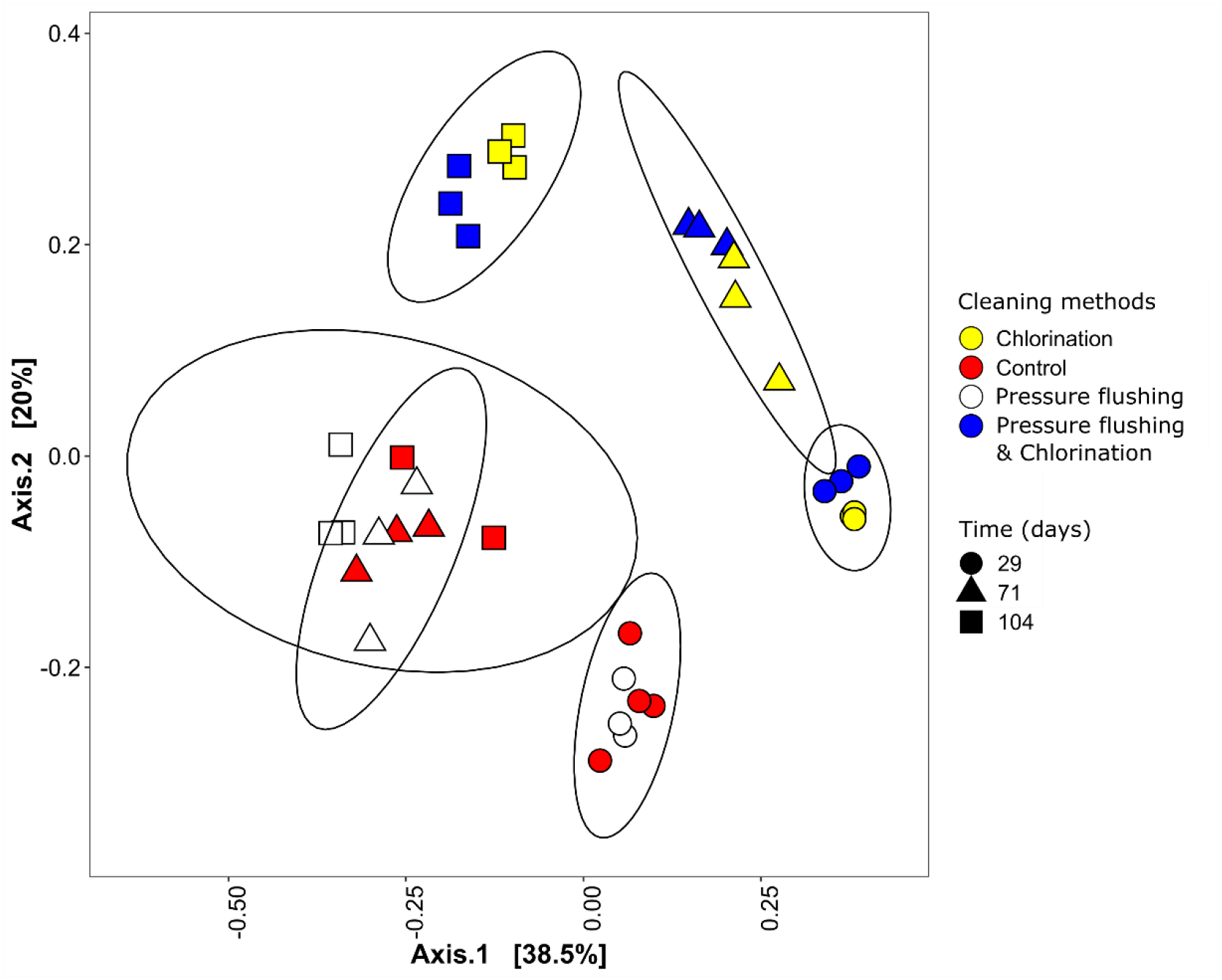
Principal coordinate analysis (PCoA) of microbial communities from dripper biofilms at genus level. The ellipsoids represent the 95% cut off similarity levels among samples.

#### Composition of bacterial communities in RWW and dripper biofilms

The biofilms and RWW microbiomes were both dominated by Proteobacteria (mainly β- and γ-Proteobacteria), Bacteroidetes, Actinobacteria and Firmicutes (Figure 8A and 8B). However, some phyla were more abundant in RWW and others more specific to biofilms. Acidobacteria were mainly detected in samples of dripper biofilms whereas Actinobacteria and Verrucomicrobia phyla were more abundant in RWW samples. In addition, the relative abundance of the Spirochaetes phylum increased over time in the dripper biofilms but was less than 1% of the overall RWW bacterial community. This means that the dripper environment induced a bacterial selection. From 70 days onwards the Synergistetes phylum appeared in RWW, explaining its increase in two drippers sampled at 104 days. This means that the dynamic of the bacterial composition of RWW has an effect on the bacterial community of dripper biofilms.

**Figure 8.**
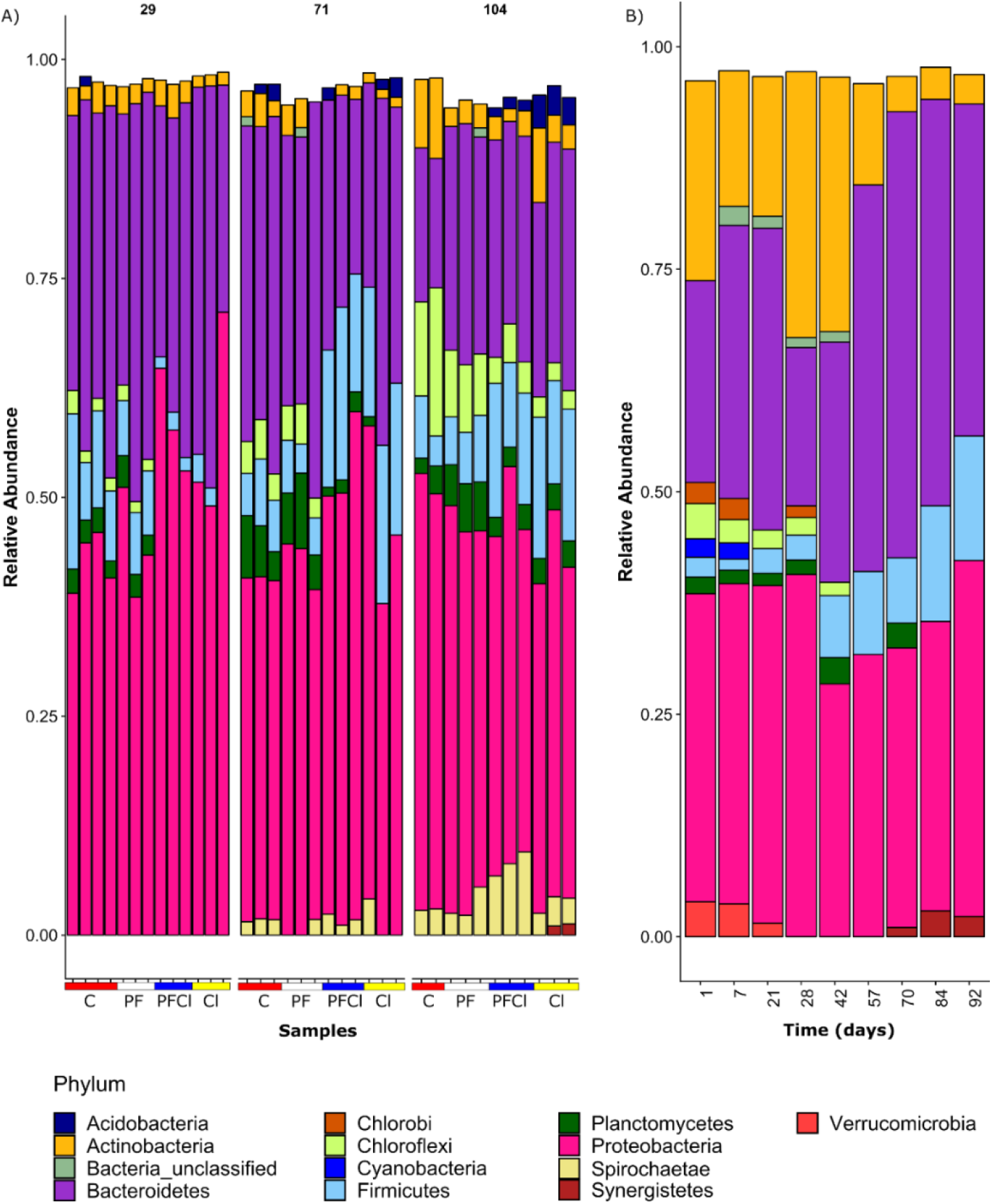
Relative abundance of bacterial phyla (>1%) in dripper biofilms over time (A) and in reclaimed wastewater (B).

Both the chlorination and pressure flushing methods modified the bacterial communities structure and composition. The relative abundance of Proteobacteria phylum was higher in chlorinated biofilms over time (Kruskal-test, p<0.05) while the relative abundance of Chloroflexi and Planctomycetes was lower (Kruskal-test, p < 0.05). Indeed, Chloroflexi and Planctomycetes phyla were under-represented (<1% of the global community) in the biofilms of the chlorinated lines compared to the non-chlorinated biofilms during the cleaning period (Figure 8A). These phyla were back in the 104 days samples once the treatment was stopped, but were still less dominant than in the non-chlorinated lines (mean relative abundances: 14% and 7% in C and PF respectively against 2% and 4% in Cl and PFCl respectively). This means that chlorination induced a bacterial selection and, although sensitive bacteria were able to grow again in the biofilm, the effect of chlorination on the communities remained 1 month after the treatments were stopped. Thus, the bacterial communities of the biofilms were driven by two factors: the dynamics of the bacterial composition of the RWW and the selection of some adapted taxa able to deal with chlorination.

As suggested with Figure 7, bacterial composition from the C and PF dripper biofilms was similar and were dominated by the same genera as members of Comamonadaceae family, *Dechloromonas*, env.OPS_17 group at 29 days and *Terrimonas*, *Dechloromonas* and *Denitratisoma* at 71 days (Table S1). In addition, the SIMPER analysis showed that there were very few genera that induced differences between the structure of C and PF communities (Figure S8), which confirmed that the bacterial structure and composition were similar between C and PF dripper biofilms. There were also few differences between the Cl and PFCl conditions (Table S1), indicating that chlorination drove the bacterial composition, as suggested in Figure 7.

The phyla whose abundance was most affected by the treatments were studied at family and genus levels in the dripper biofilms. Proteobacteria were mainly composed of β-Proteobacteria, followed by γ-Proteobacteria, δ-Proteobacteria and α-Proteobacteria. Relative abundances of β-Proteobacteria and α-Proteobacteria were statistically higher in chlorinated biofilms (Cl and PFCl, Kruskal test, p < 0.05) while γ-Proteobacteria were not impacted by the cleaning method used (Kruskal test, p > 0.05). Inversely, δ-Proteobacteria were statistically higher in non-chlorinated drippers (C and PF) with 2% against <1% in chlorinated drippers (Cl and PFCl). Among the β-Proteobacteria and α-Proteobacteria, the Comamonadaceae member’s family, *Azospira,* Sphingomonadaceae, *Caulobacter* and *Hyphomicrobium* (α-Proteobacteria) were more abundant in chlorinated biofilms (Kruskal test, p<0.05) whereas others were more abundant in the C and PF as *Denitratisoma* and *Aquabacterium* (Kruskal test, p<0.05) (Figure 9, Table S1). SIMPER analysis indicated that *Azospira* and an unclassified genus from *Comamonadaceae* were the main genera responsible for the clustering of the bacterial community depending on the use of chlorination (Figure S8) during the cleaning period. The genus *Pseudoxanthomonas* (γ-Proteobacteria) were more abundant in Cl and PFCl (3-6%) conditions compared to C and PF conditions (<1%) whereas the Run-SP154_ge was more represented in non-chlorinated drippers (C and PF).

**Figure 9.**
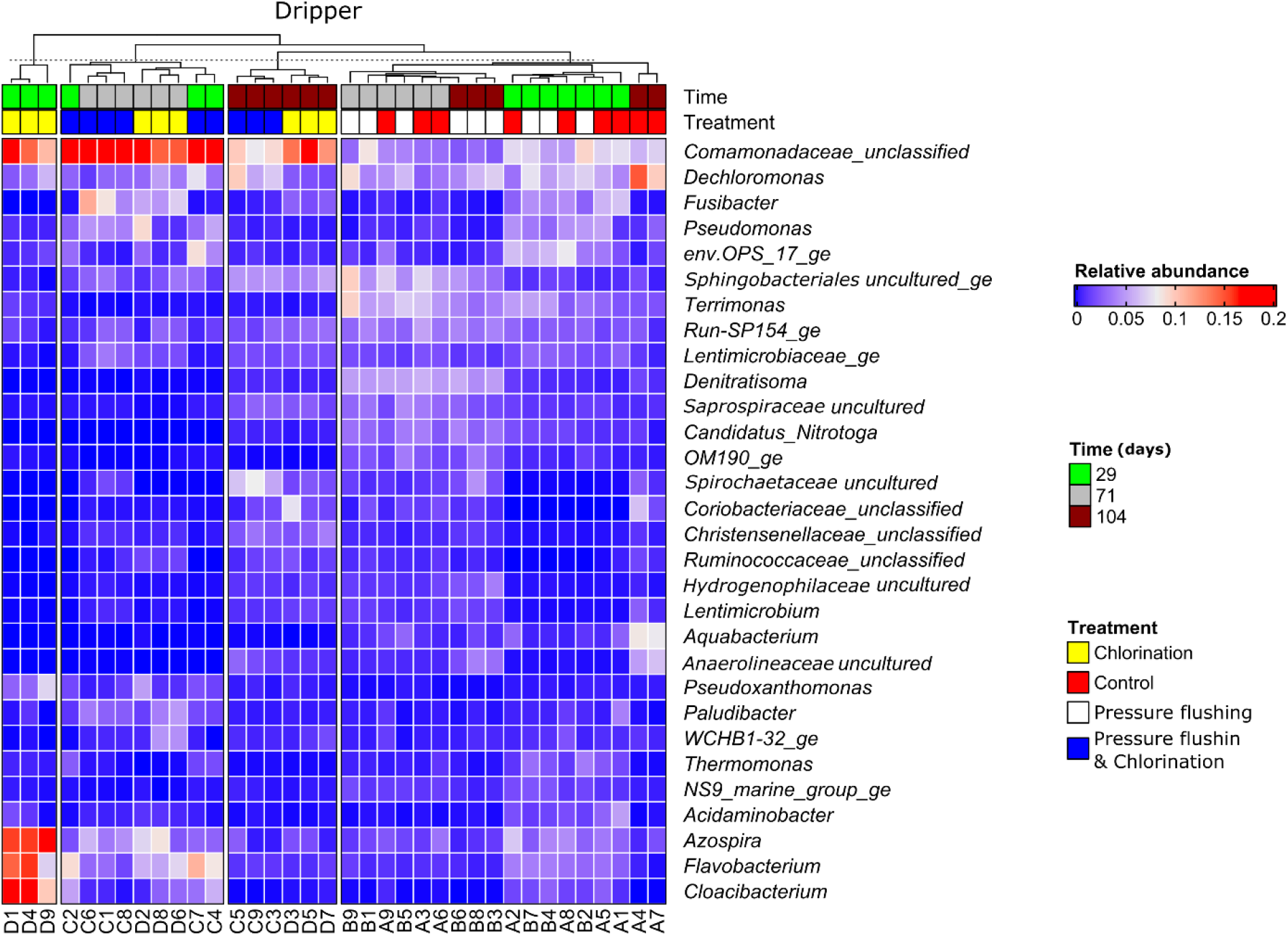
Heat map of bacterial genera from dripper biofilms. Genera with the top 30 relative abundance are shown.

The relative abundance of Firmicutes (dominated by Clostridia class) were significantly higher in chlorinated biofilm compared to non–chlorinated biofilm (Kruskal-test, p<0.05) at 71 days (4-6% in C-PF against 16-17% in Cl and PFCl respectively, Figure 8A). Moreover, *Fusibacter* genus (Clostridia class) drove the separation of the bacterial community structure according to the use of chlorine (Figure S8).

Chloroflexi (mainly composed of one Anaerolineaceae member’s family, *Leptolinea*, Choroflexi SBR2076 and SJA-15 group) and Planctomycetes phylum (mainly composed by OM190 group, *Planctomyces* and *Schlesneria* genera) were significantly lower in the chlorinated biofilms (Cl and PFCl) during the treatment period (Figure 8A).

#### Composition of eucaryotic communities in dripper biofilms

At the end of the cleaning period, the 18S rRNA sequence analysis also showed an effect of the treatments on the structure and composition of the eukaryotic communities (Figure S9). The Ciliophora and Fungi classes were the most represented. Among the Ciliophora, the family Tokophryidae was mainly present in the biofilms of non-chlorinated drippers, while the families Vorticellidae and Peritrichia were mainly present in the biofilms of chlorinated drippers. The fungi, dominated by the families Dothideomycetes or Pezizomycotina, were lower in chlorinated drippers whereas the Sordariomycetes family was mainly present in the chlorinated biofilms.

#### Impact of the interruption of the cleaning method on the bacterial community composition

SIMPER analysis at 104 days showed that there was no difference between C and the other conditions (Table S8), which confirmed that bacterial communities converged (Figure 7) when the treatments were stopped.

Chloroflexi and Planctomycetes phyla increased in the biofilms of the chlorinated drippers from Cl and PFCl lines when chlorination was stopped (Figure 8A). The genera and taxonomic groups belonging to Chloroflexi such as the Anaerolineaceae member’s family, *Leptolinea*, SBR2076 and SJA-15groups settled in chlorinated dripper biofilms. Other minor bacterial families in chlorinated drippers during the cleaning period re-emerged such as Saprospiraceae (Bacteroidetes), Spirochaetaceae (Spirochaetae), Christensenellaceae (Firmicutes) and Hydrogenophilaceae (β-Proteobacteria) at levels similar to biofilms from non-chlorinated drippers (Figure 9). Inversely, the relative abundance of the main genera from chlorinated biofilm during the cleaning period decreased for *Azospira*, *Flavobacterium* and *Cloacibacterium* (Figure 9).

A shift was observed between the bacterial compositions from the C drippers at 104 days compared to 71 days mainly due to the increase of the relative abundances of Chloroflexi (from 3% to 14%) and Actinobacteria (from 3 to 8.5%), which were higher than in the other conditions (p-value < 0.05). On the contrary, the relative abundance of Bacteroidetes phylum (represented by *Terrimonas* genus) decreased from 37% to 16%. The dominant genera already found during the cleaning period increased such as *Dechloromonas* and *Aquabacterium* (β-Proteobacteria) (p-value > 0.05), which could explain the decrease of 1/Simpson’s index observed (Table 4). The relative abundances of Chloroflexi, mainly composed of Anaerolineaceae member’s family, and Actinobacteria were higher in the C condition than in the others.

## 4. Discussion

In this study, the effect on biological clogging and microbial communities of different cleaning methods (chlorination (Cl), pressure flushing (PF), and a combination of both methods (PFCl)) was investigated. Commercial flat drippers (1 l.h^−1^) used in agriculture were installed on a test bench and supplied with reclaimed wastewater. After a 2-month period of treatments, cleaning events were interrupted in order to study the impact on biofouling regrowth and bacterial communities.

### Chlorination but not purge-only treatment helped maintain flow rates

The effectiveness of cleaning methods is often looked at through flow maintenance (Feng et al., 2018; Han et al., 2018; Song et al., 2017). Chlorination (1.5 ppm) used alone or in combination with pressure flushing (Cl and PFCl), maintained flow rates and reduced distribution heterogeneity throughout the cleaning period. In contrast, the uniformity of water distribution decreased over time under non-chlorinated conditions (C and PF), meaning that using chlorination is preferable to maintain the expected flow rates. Chlorination results were consistent with those of J. Li et al. (2010) who showed that chlorinating to 2.5 ppm free chlorine once a week maintained the expected dripper flow rates supplied by reclaimed wastewater for a longer period of time. However, the impact of the pressure flushing is not consistent with Yu et al., (2018) who showed that increasing the pressure from 0.1MPa to 0.3MPa for 2 minutes helped maintain the expected flow rates longer by removing large particles (< 6.6 μm in diameter). However, the biofilm is considered to be an elastically deformable but quasi-static structure, allowing it to deal with an increase of pressure and flow velocity (Picioreanu et al., 2018). Moreover, biofilms can increase the pressure loss due to friction along the labyrinth (Li et al., 2016): the biofilm formed in the labyrinth increases roughness, which in turn increases friction, resulting in increased flow resistance and thus increased the pressure loss with a consequent decrease in the uniformity of the distribution. Thus, in the case of the pressure flushing condition, the increase in pressure may not have been sufficient to promote a detachment of biofilm and maintain flow rates.

### Impact of the cleaning procedures on biofouling

Purging had no impact on bacterial concentration and biofouling (volume, thickness). A recent study showed that flushing (1h per day) with reclaimed wastewater is not effective in limiting the clogging of drippers compared to flushing with fresh water due to the presence of suspended solids and other organic compounds (N. Li et al., 2019). Thus, the use of freshwater could be a solution to improve the efficiency of the pressure flushing method. However in the case of field irrigation with treated wastewater it is not always possible to have access to a second water resource.

Conversely, the level of biofouling and bacterial concentration in the chlorinated lines (Cl and PFCl), mainly in the PFCl line, tended to decrease compared to non-chlorinated lines. The combination of pressure flushing and chlorination seemed to better control biofouling than chlorination alone. This decrease was mainly due to a significant decrease in biofouling in the middle of the channel, where the flow velocity was higher (Ait-Mouheb et al., 2018; Al-Muhammad et al., 2016; Yu et al., 2018a). Increasing the flow velocity by increasing the pressure promotes mass transfer (Beyenal and Lewandowski, 2002; Moreira et al., 2013) and influences mixing (Khaydarov et al., 2018; Naher et al., 2011). Mathieu et al. (2014) showed in a rotating disc reactor that the shear stress and cohesive force necessary to detach the biofilm decreased after chlorine application (10 ppm during 1h). This is probably due to breaking the hydrogen bond, polymer and hydrophobic interactions caused by chlorine (Xue and Seo, 2013), which weakens the biofilm and makes it easier to detach. Thus, chlorine mass transfer caused by hydrodynamic conditions in the main flow zones (middle of the milli-channel) may explain why biofouling level was lower under PFCl conditions than in the C and PF conditions in the middle and corners of the channel and why bacterial concentration was lower in this condition. However, the combination of the two methods did not completely suppress the development of biofouling. One explanation is that the increase in the PFCl condition favours the compaction of the biofilm and the resistance of the basal layer in the central and corner areas (Blauert et al., 2015; Rochex et al., 2008; Valladares Linares et al., 2016; Wagner and Horn, 2017), which reduced the chlorine effect (Lee et al., 2018). Lee et al. (2018) showed that the free chlorine penetration at 10 ppm of Cl_2_ was low in a biofilm of 2000 μm (7 days to reach the substratum) and tended to promote the sloughing of the superficial layers of the biofilm. Thus, the concentration of free chlorine used (1.5 ppm) may be insufficient to reduce the level of clogging, especially in corner areas where the thickness was higher.

### Chlorination strongly modified microbial communities in biofilms

In this study, total bacteria, diversity and richness index increased with the level of clogging. The bacterial diversity was lower in chlorinated conditions compared to the control and pressure flushing line. The diversity of the biofilms of drinking water distribution pipes was also previously shown to decrease with the increasing concentration of chlorine (0.05 to 1.76 ppm) (Fish and Boxall, 2018; Mi et al., 2015).

No effect of the pressure flushing on the bacterial community was found in comparison with the non-treated control line, which is consistent with Li et al. (2015) who studied the microbial composition of biofilms by PLFAs analysis. On the other hand, the use of chlorine, combined or not with pressure flushing, has significantly altered the structure of the bacterial community during the cleaning step. Proteobacteria and Firmicutes phyla appeared to contain bacterial members resistant to chlorine treatment whereas Chloroflexi phylum appeared to be sensitive to chlorine which is consistent with Song et al. (2019). Chloroflexi phylum includes many filamentous bacteria such as Anaerolineaceae member’s family and *Leptolinea* genus detected in sequencing data and responsible for clogging membrane bioreactors in wastewater treatment plants (Li et al., 2008) and drippers supplied by RWW (Lequette et al., 2019). Chloroflexi may have a key role in the clogging of drippers.

Chlorinated dripper biofilms were largely dominated by OTUs belonging to the Comamonadaceae family (β-Proteobacteria), followed by *Flavobacterium* (Bacteroidetes), *Pseudomonas* and *Pseudoxanthomonas* genera (γ-Proteobacteria). β-Proteobacteria are frequently found in chlorinated drinking distribution systems and wastewater biofilms (Douterelo et al., 2016; Navarro et al., 2016; Shaw et al., 2014) and their abundance can increase with the chlorine concentration (Wang et al., 2019b). The Comamonadaceae family was commonly found in water and chlorinated biofilms (Wang et al., 2019a; Zhang et al., 2019). *Pseudomonas* and *Hydrogenophaga* (Comamonadaceae) are known to exhibit resistance to disinfection (Jia et al., 2015; Wang et al., 2019a). Increases in relative abundance of *Pseudomonas* have been observed in water samples after chlorination (Jia et al., 2015) and *Hydrogenophaga* are known to colonise chlorinated distribution pipes fed with reclaimed wastewater (Wang et al., 2019b).

Firmicutes predominated in the chlorinated drippers after 2 months of treatment. At the class level, it was mainly composed of Clostridia (abundancy between 10 and 14%), which includes many members capable of producing endospores. These endospores are highly resistant to a variety of environmental challenges, such as heat, solvents, oxidising agents, UV irradiation and desiccation (Abecasis et al., 2013). The resilience of endospores allows them to remain viable in a hostile environment for long periods of time, and contributes to their survival and proliferation in chlorinated environments (Douterelo et al., 2016).

The eukaryotic communities, mainly composed of Ciliophora and Fungi, were also influenced by the use of chlorine. Sordariomycetes, Tremellomycetes and Chytridiomycetes were still present in chlorinated biofilms, when Dothideomycetes or Pezizomycotina were only recovered in control and purge drippers. Sordariomycetes, Tremellomycetes and Chytridiomycetes are commonly found in the effluent of wastewater treatment plants and are involved in water treatment (Assress et al., 2019). Sordariomycetes has also been detected as dominant in biofilms from drinking water distribution systems (0.05 ppm to 0.8 ppm of free chlorine) (Fish and Boxall, 2018). These results suggest that among fungi, some are more tolerant to chemical disinfection. The resistance to chemical disinfection of some fungi can be explained by their thick melanized cell walls, which increases resistance to mechanical damage and limits the intrusion of biocides into the cell (Hageskal et al., 2012) or their ability to form spores (Sonigo et al., 2011). Ciliophora were affected by the use of chlorine, with chlorine-sensitive families (e.g. Tokophryidae) whereas others were still present in chlorinated drippers (e.g. Vorticellidae). Ciliophora contains predators of bacteria and can significantly reduce the concentration of the bacteria (Parry et al., 2007) and influences the morphology of biofilms (Böhme et al., 2009; Derlon et al., 2012). Although eukaryotic communities from dripper biofilms are poorly studied, their role in biofilm development and resistance to cleaning processes may be important, and must be integrated in studies on the control of biofilms.

### Resilience of microbial communities once treatments are stopped

After 1 month without cleaning events, several bacterial families detected as chlorine-sensitive re-emerged such as Saprospiraceae, Spirochaetaceae, Christensenellaceae and Hydrogenophilaceae at levels similar to biofilms from non-chlorinated drippers. The relative abundance of Chloroflexi phylum also increased significantly after this period. The concentration of bacteria and diversity and richness indexes also increased. This indicates that stopping treatments reduces selection pressure and makes it easier to recruit new species. Fish and Boxall (2018) have found a similar result in the case of biofilms developing in drinking water distribution systems and this would explain the increase of clogging of drippers after the interruption of chlorination observed by Katz et al. (2014) (10 ppm of free chlorine). This also illustrates the “resilient” ability of the bacterial community to deal with the chlorine used (Allison and Martiny, 2009) in a drip irrigation system. However, PCoA analyses showed that the drippers in the chlorinated lines still form a different cluster than non-chlorinated lines 1 month after stopping the treatment, showing that the history of the treatment still affects the structure of the communities and that the resilience was only partial.

## 5. Conclusion

The combined use of OCT and high throughput sequencing highlighted the impact of the two cleaning procedures (pressure flushing, chlorination) on biofouling of drip irrigation systems supplied by RWW:

– The pressure flushing method alone did not reduce the biofouling of the drippers and did not influence the microbiome of the biofilm.
– On the other hand, chlorination application alone or combined with pressure flushing steps reduced biofouling of the drippers, mainly in the main flow zone.
– Some bacteria commonly found to clog systems such as Chloroflexi and Planctomycetes appear to be sensitive to chlorine and regrowth after chlorination is discontinued.
– Inversely, others are more resistant such as the members of the Comamonadaceae and Clostridia (Firmicutes). Further research into the mechanisms of resistance is needed to improve the control of biofilms.
– The suspension of treatment leads to an increase in the bacterial diversity of the initially chlorinated biofilms and a convergence of the communities towards non-chlorinated biofilms, although differences in the structure remains. This indicates that the bacterial community is resilient to chlorine, but that the community at a global scale keeps the ‘memory’ of the chlorination.

As indicated, the concentration of chlorine used may have been a determining factor, especially in low-velocity vortex areas where the thickness of the biofouling was higher. Further studies on chlorine transfer, whether or not combined with purging, would allow a better understanding of the mechanisms of biofilm maintenance in these systems. In addition, the microbiome of reclaimed water evolved in time and depended on the treatment used. For this reason, additional studies to investigate the effect of the wastewater treatment (i.e. activated sludge, membrane bioreactor) as well as the role of bacterial predators on the microbial composition of a biofilm, have to be performed.

## 6. Conclusion

The combined use of OCT and high throughput sequencing highlighted the impact of cleaning procedures (pressure flushing, chlorination) on biofouling of drip irrigation systems supplied by RWW:

– Pressure flushing method alone did not reduce the biofouling of the drippers and did not influence the microbiome of the biofilm.
– On the contrary, chlorination application alone or combined with pressure flushing step reduced biofouling of the dripper, mainly in the main flow zone.
– Some bacteria commonly found in clogging systems such as Chloroflexi and Planctomycetes appear to be sensitive to chlorine but re-grow after chlorination is discontinued.
– Inversely, others are more resistant such as Comamonadaceae member’s family and Clostridia (Firmicutes). Further research into the mechanisms of resistance would improve the control of biofilms.
– The suspension of treatments leads to an increase in the bacterial diversity of the initially chlorinated biofilms and a convergence of the communities towards non-chlorinated biofilms, although differences in the structure remains. This indicates that the bacterial community is resilient to chlorine, but that the community at a global scale keep the ‘memory’ of the chlorination.

As indicated, the concentration of chlorine used may have been a determining factor, especially in low velocity vortex areas where the thickness of the biofouling was higher. Further studies on chlorine transfer, whether or not combined with purging, would allow a better understanding of the mechanisms of biofilm maintenance in these systems. In addition, the microbiome of reclaimed water evolved in time and depends on the treatment used. For this reason, additional studies to investigate the effect of the wastewater treatment (ie activated sludge, membrane bioreactor) as well as the role of bacterial predators on the microbial composition of a biofilm, have to be performed.

## Acknowledgements

The authors gratefully acknowledge the financial support of the French Water Agency, project “Experimental platform for the reuse of reclaimed wastewater in irrigation, Murviel-lès-Montpellier” (Project 2017-1399). We thank Guillaume Guizard (LBE-INRAE, Narbonne, France) and Jean-François Bonicel (UMR ITAP-INRAE, Montpellier, France) for their contribution to the development of the dripper system and Annabelle Mange (UMR G-Eau-INRAE, Montpellier, France) for their assistance.

## Supplementary data

**Figure S1.**
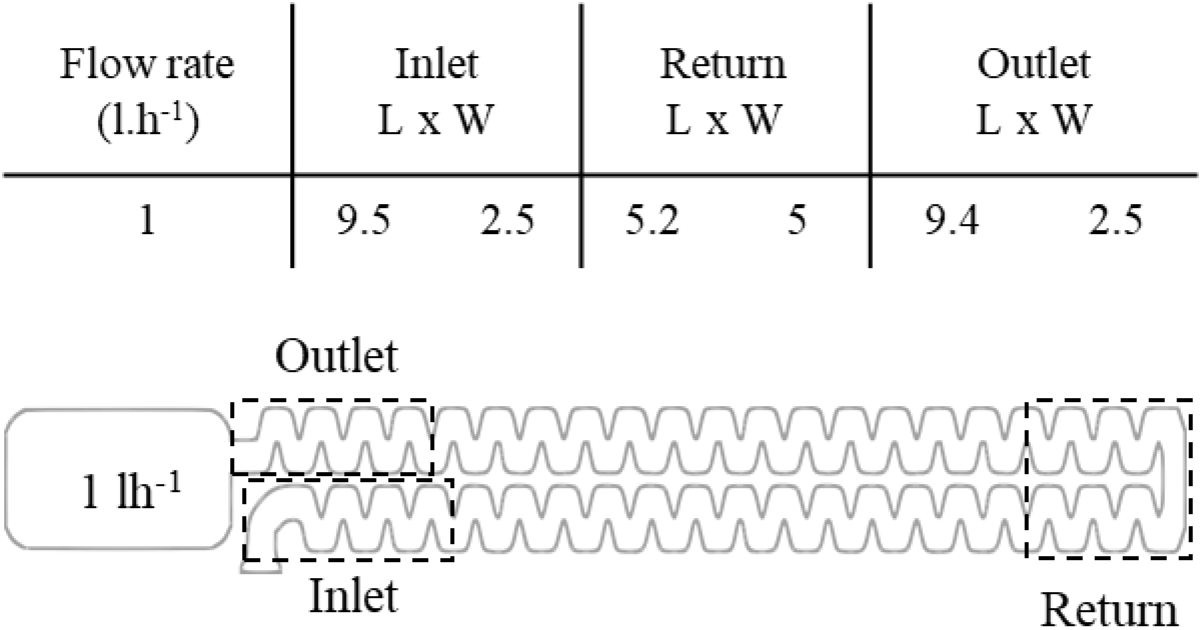
Measurements of the different areas of the labyrinth. L: length and W: width.

**Figure S2.**
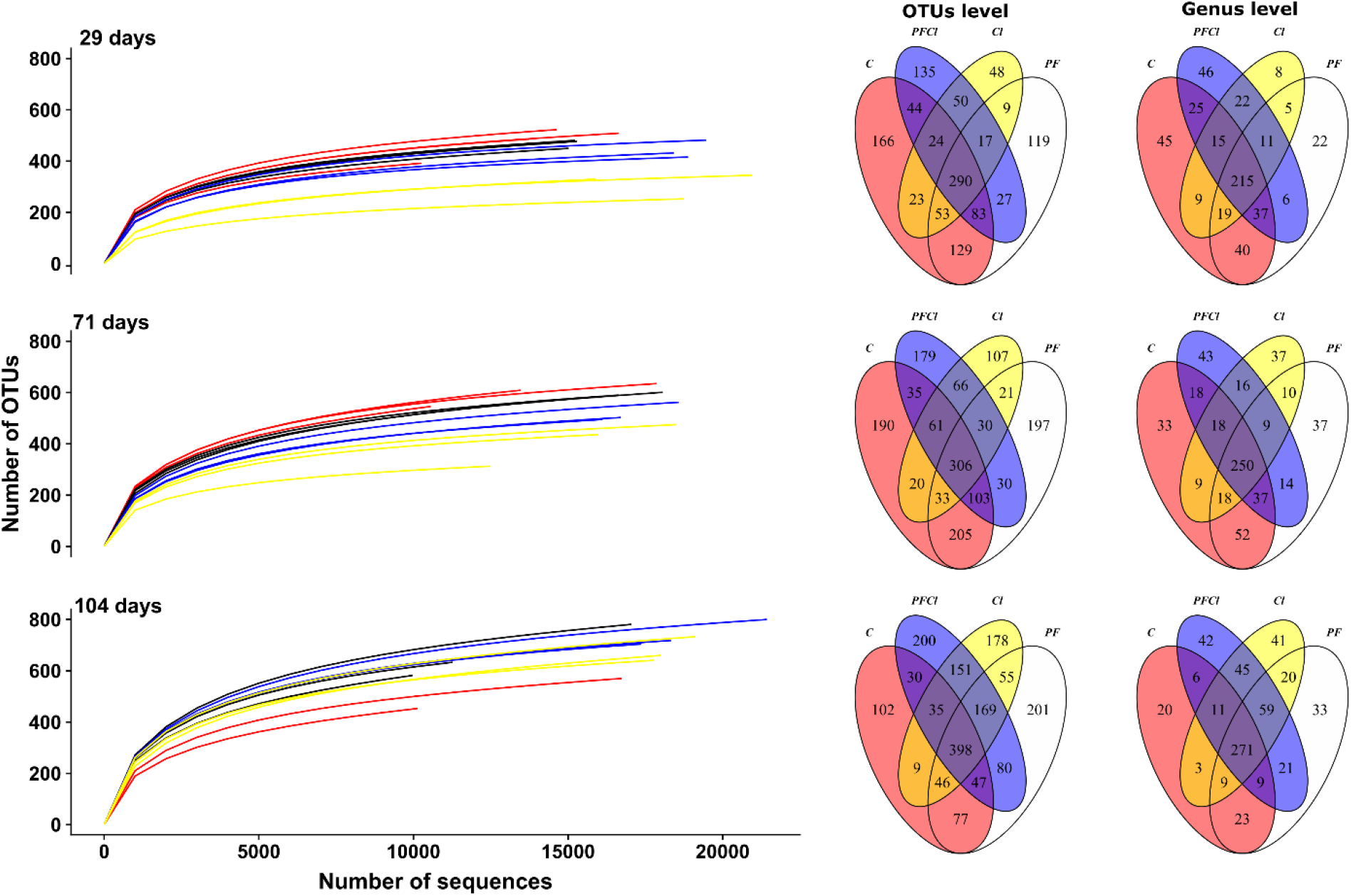
Rarefaction curves from dripper biofilms and Venn Diagram of bacterial OTUs and bacterial genera in time. (Control: red, Pressure flushing: white, Pressure flushing & Chlorination: blue, Chlorination: yellow).

**Figure S3.**
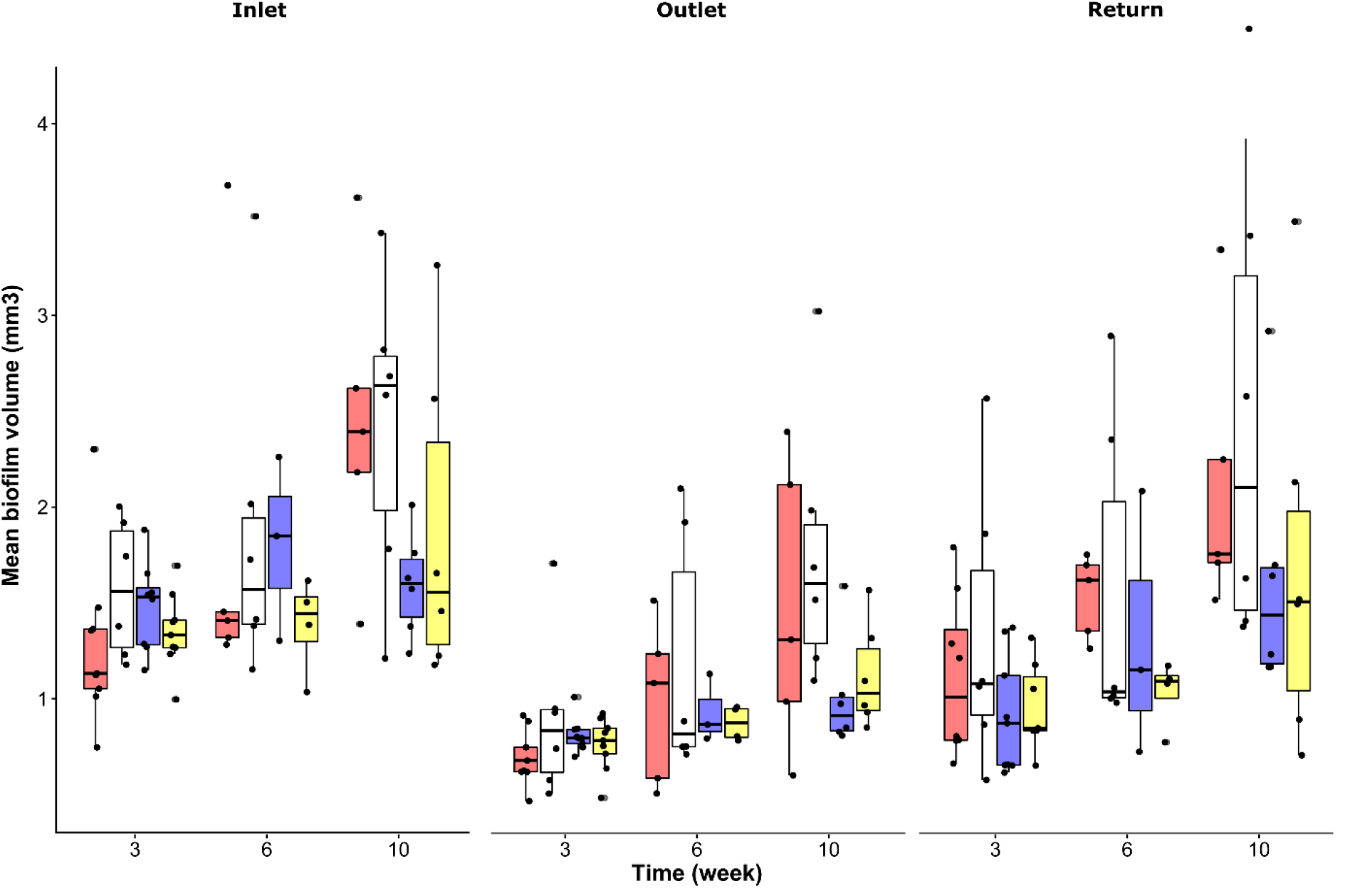
Biofilm volume during the cleaning period. Control 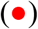, Pressure flushing (○), Chlorination 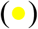 and Pressure flushing/Chlorination 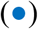.

**Figure S4.**
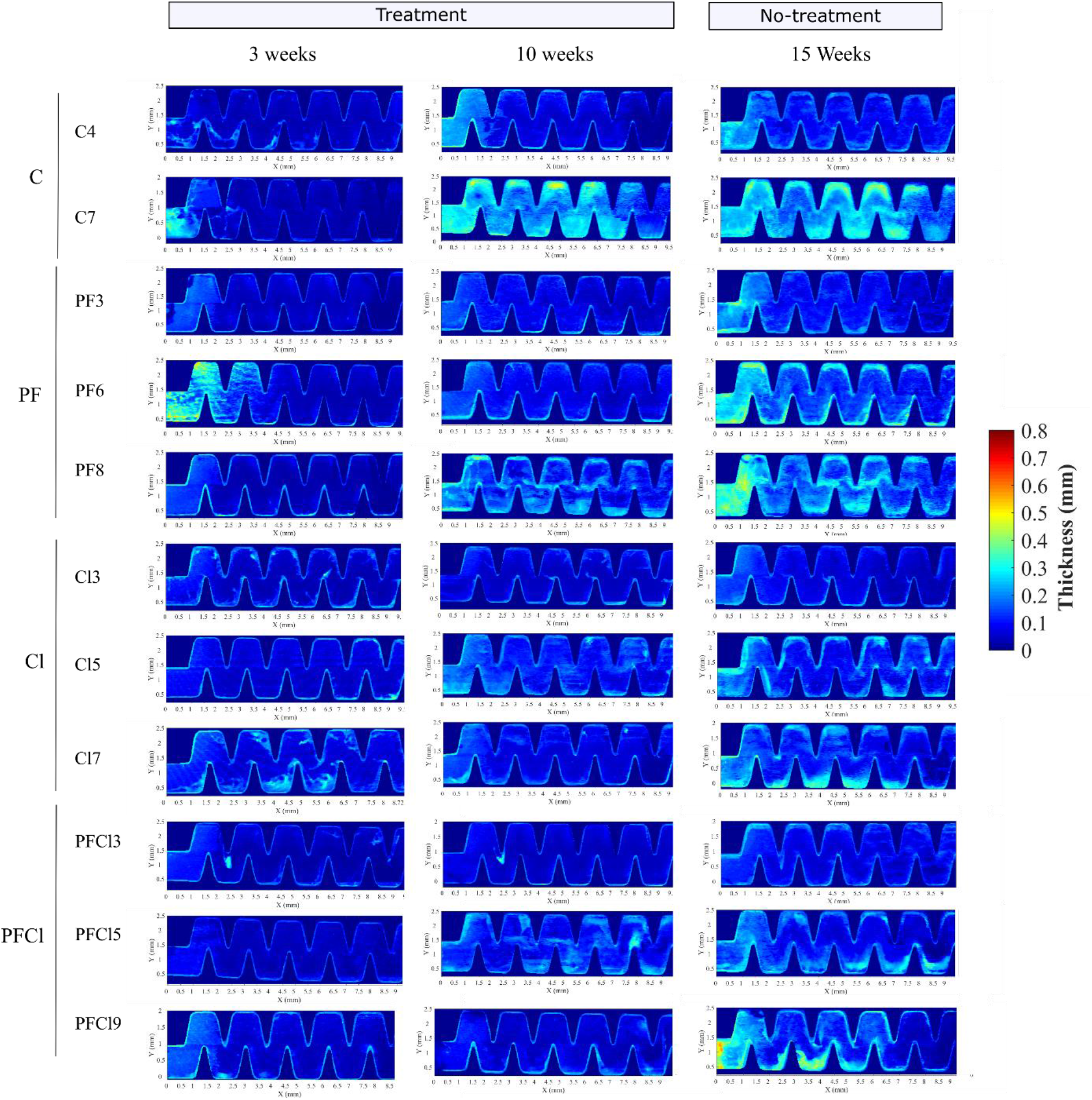
Biofilm thickness at the outlet of drippers under the C (Control), PF (Pressure flushing), Cl (Chlorination) and PFCl (Pressure flushing/Chlorination) conditions measured at 3, 10 and 15 weeks. The drippers presented are those also analysed by 16S rRNA sequencing at 104 days.

**Figure S5.**
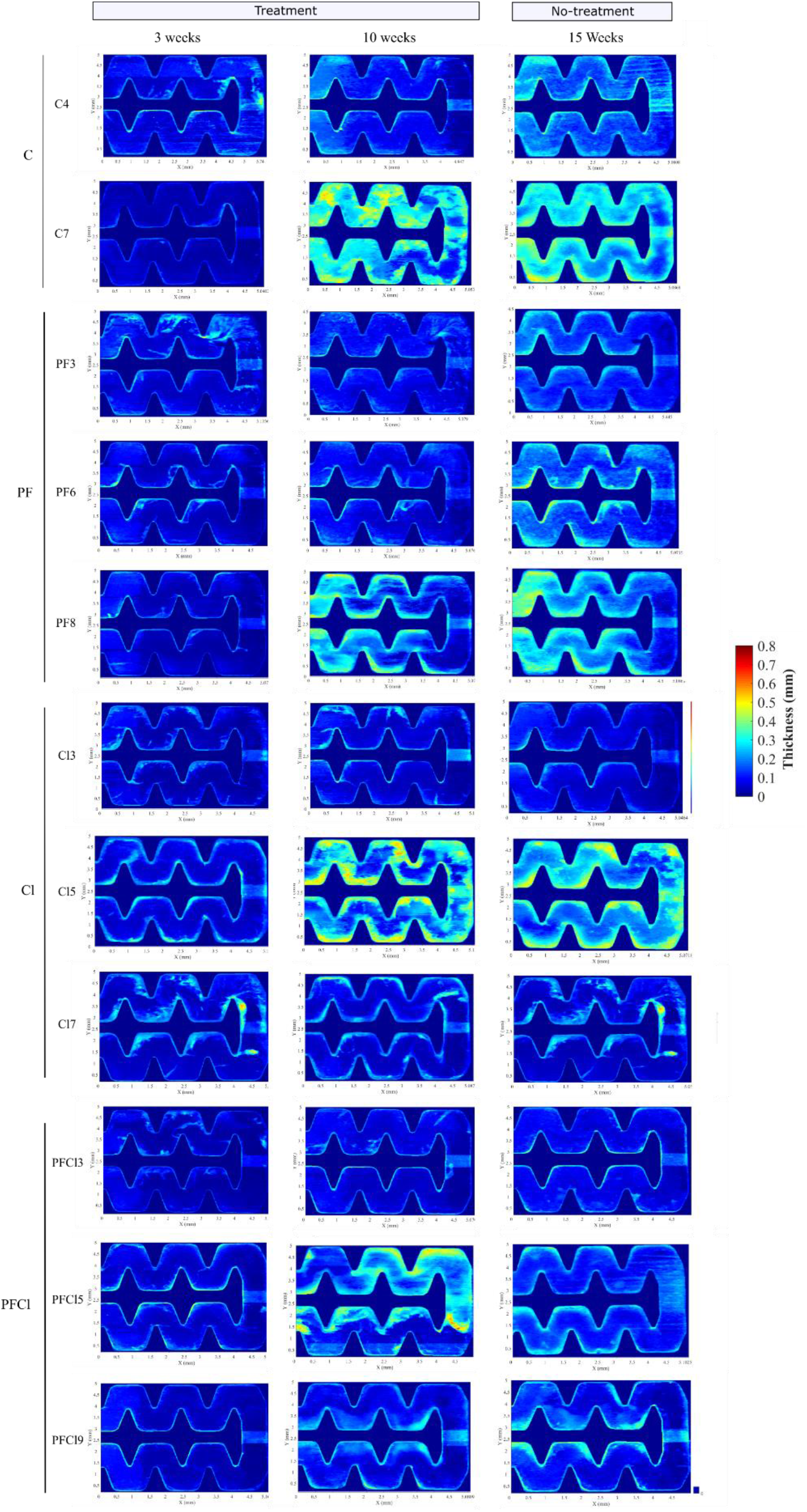
Biofilm thickness at the return of drippers under the C (Control), PF (Pressure flushing), Cl (Chlorination) and PFCl (Pressure flushing/Chlorination) conditions measured at 3, 10 and 15 weeks. The drippers presented are those also analysed by 16S rRNA sequencing at 104 days.

**Figure S6.**
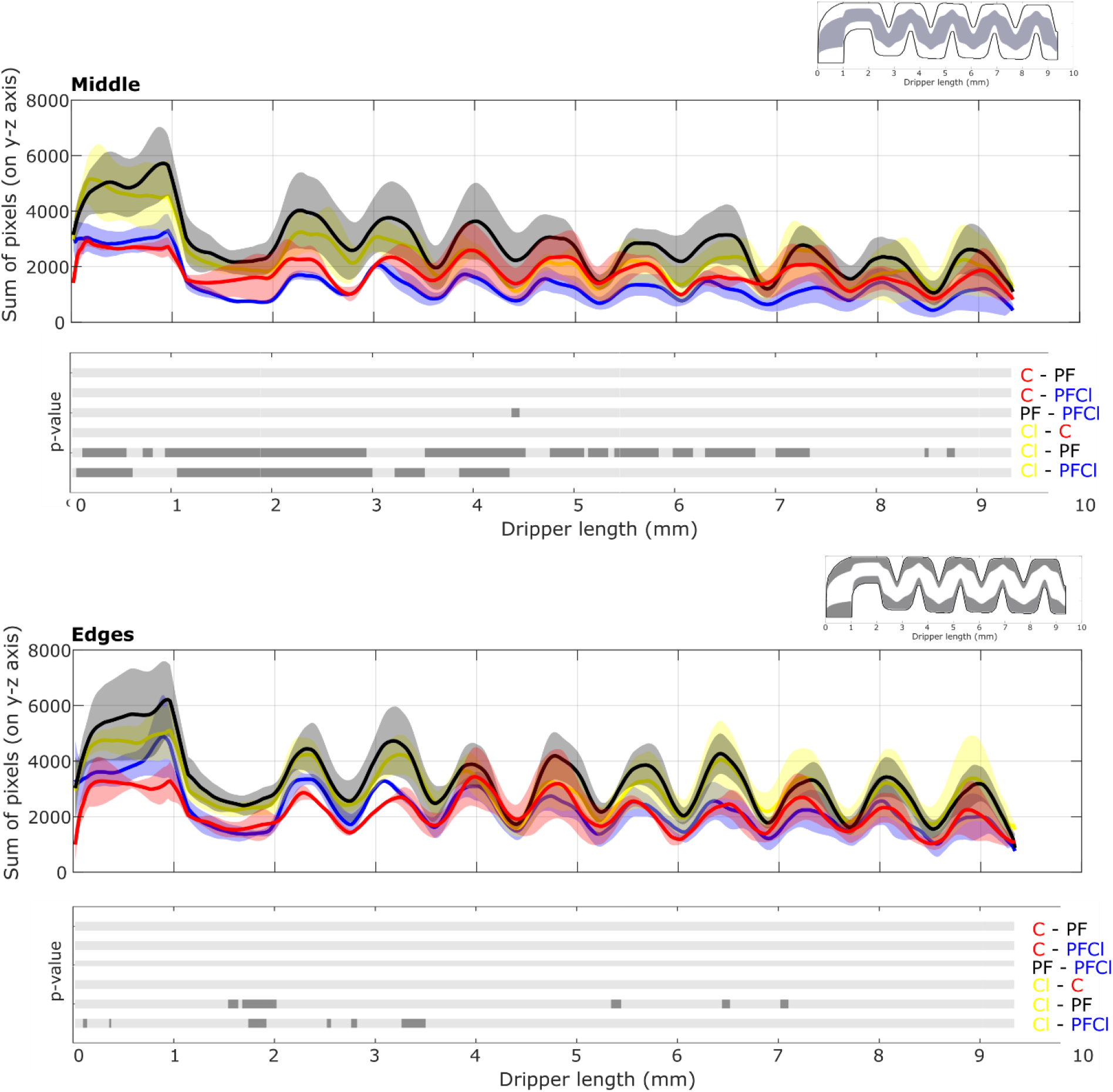
Mean number of pixels associated with the biofilm mass (and standard deviation) in the middle and edges of the inlet dripper channel after one month without cleaning. Control (C-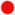), Pressure flushing (PF-●), Chlorination (Cl-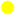) and Pressure flushing combined with Chlorination (PFCl-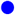); n=6 per condition. P-value graphs show the results of the Wilcoxon tests with 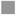: non-significant, 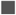: p<0.1, ■: p<0.05.

**Table S1.**
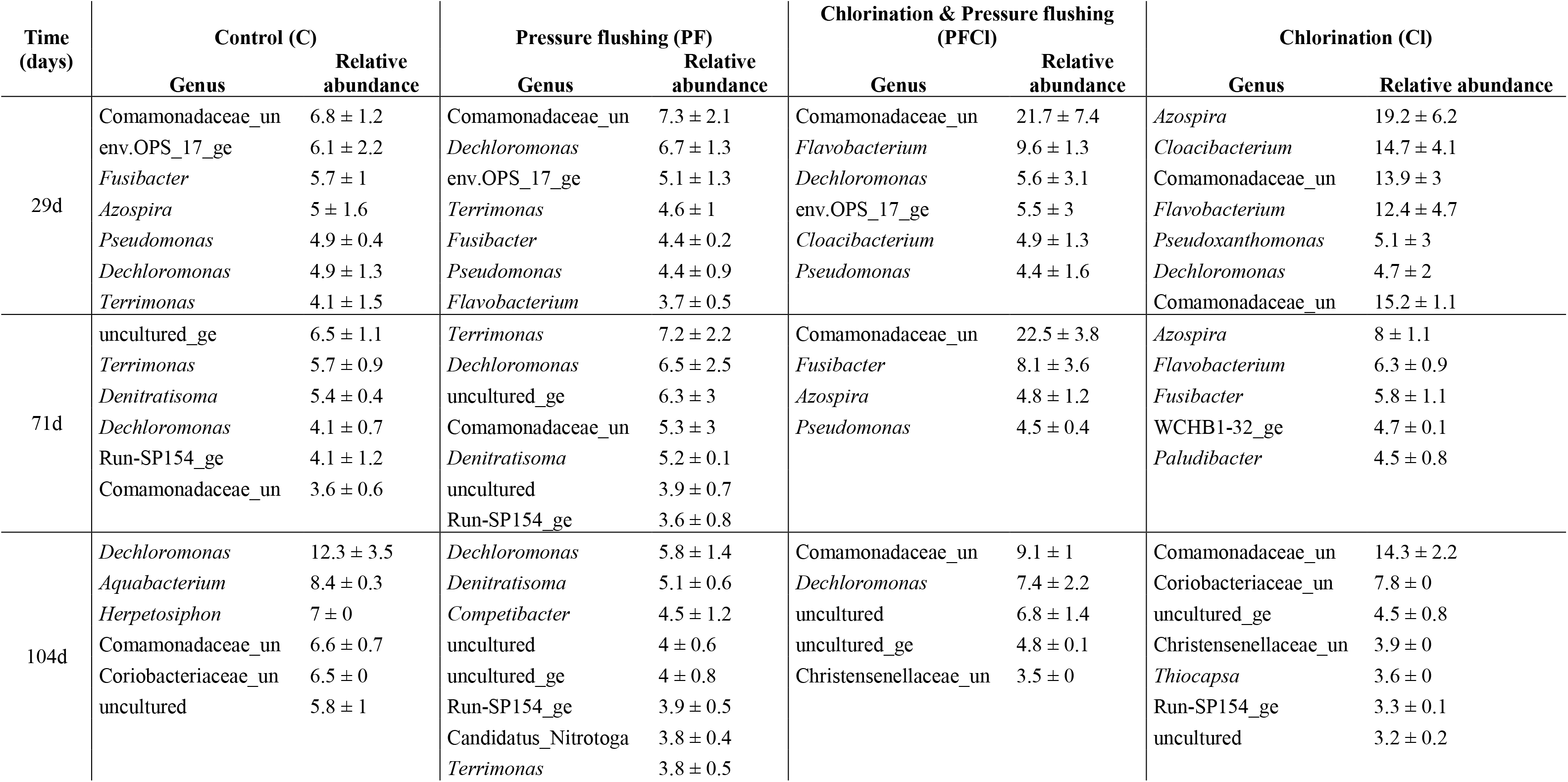
Top abundant genera (<3%, ×10^−1^) of genera found specifically in one dripper type for each sampling time.-un: unclassified, _ge: genus.

**Figure S8.**
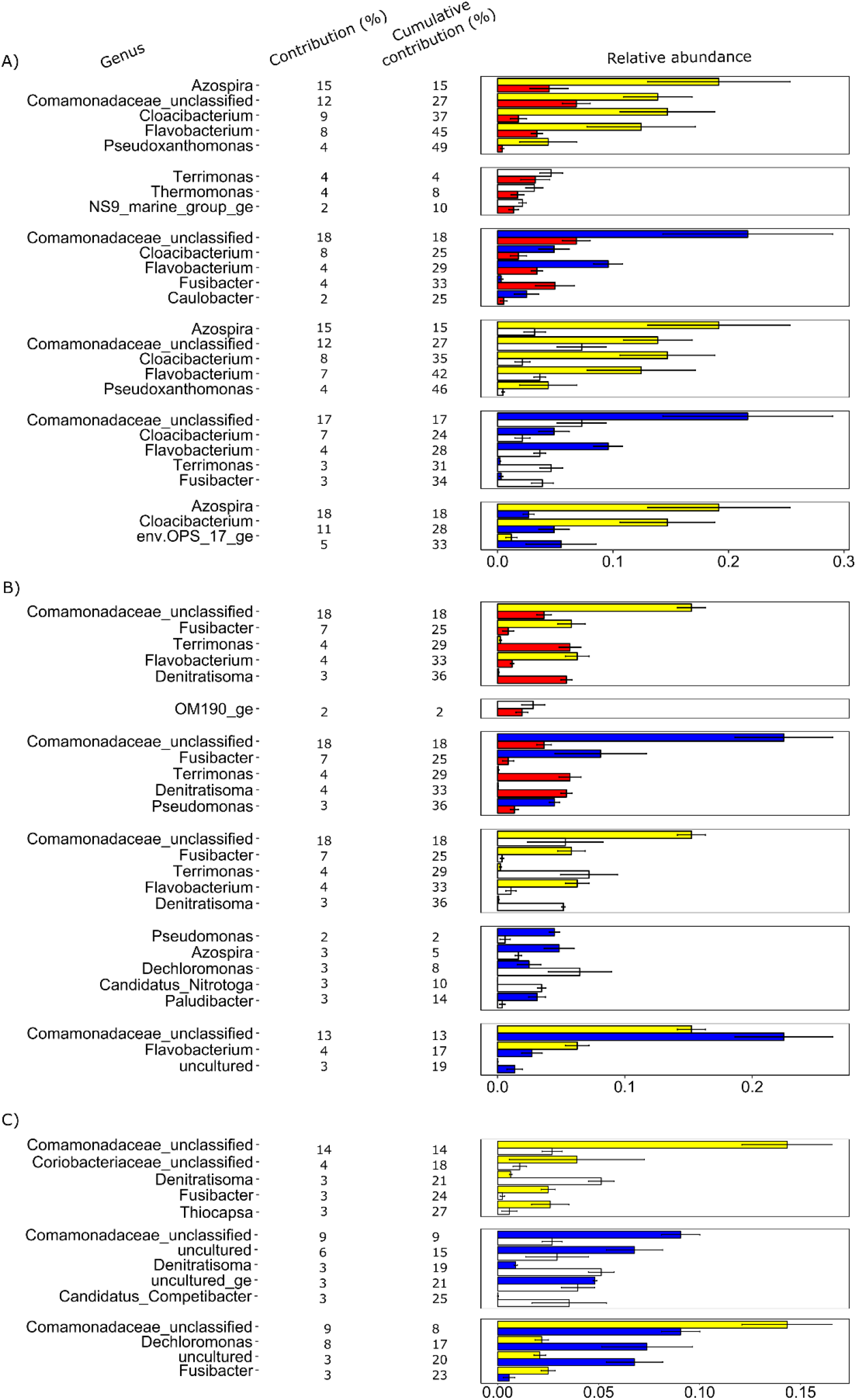
Most influential bacterial genera in discriminating between 2 conditions (SIMPER analysis). Control 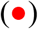, Pressure flushing (○), Chlorination 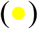 and Pressure flushing/Chlorination 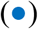.

**Figure S9.**
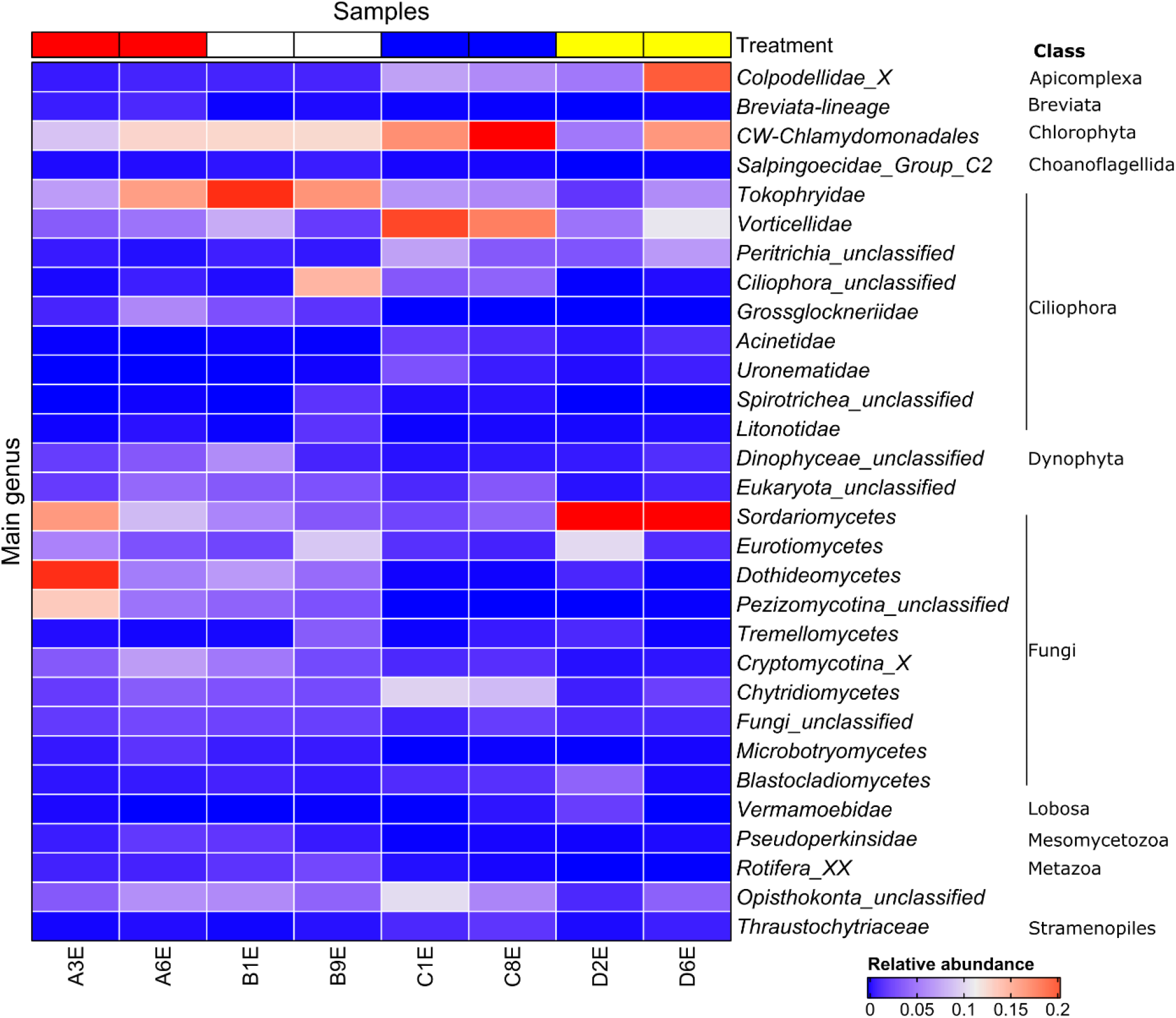
Heat map of eukaryotes genera from dripper biofilms. Genera in top thirty relative abundance are shown. Control 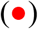, Pressure flushing (○), Chlorination 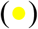 and Pressure flushing/Chlorination 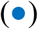.

