## Supplemental for "Effect of chlorination and pressure flushing of drippers fed by reclaimed wastewater on biofouling"

### Supplementary data

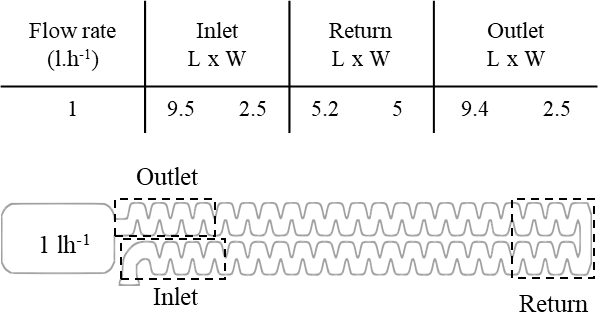

**Figure S1 Measurements of the different areas of the labyrinth. L: length and W: width**

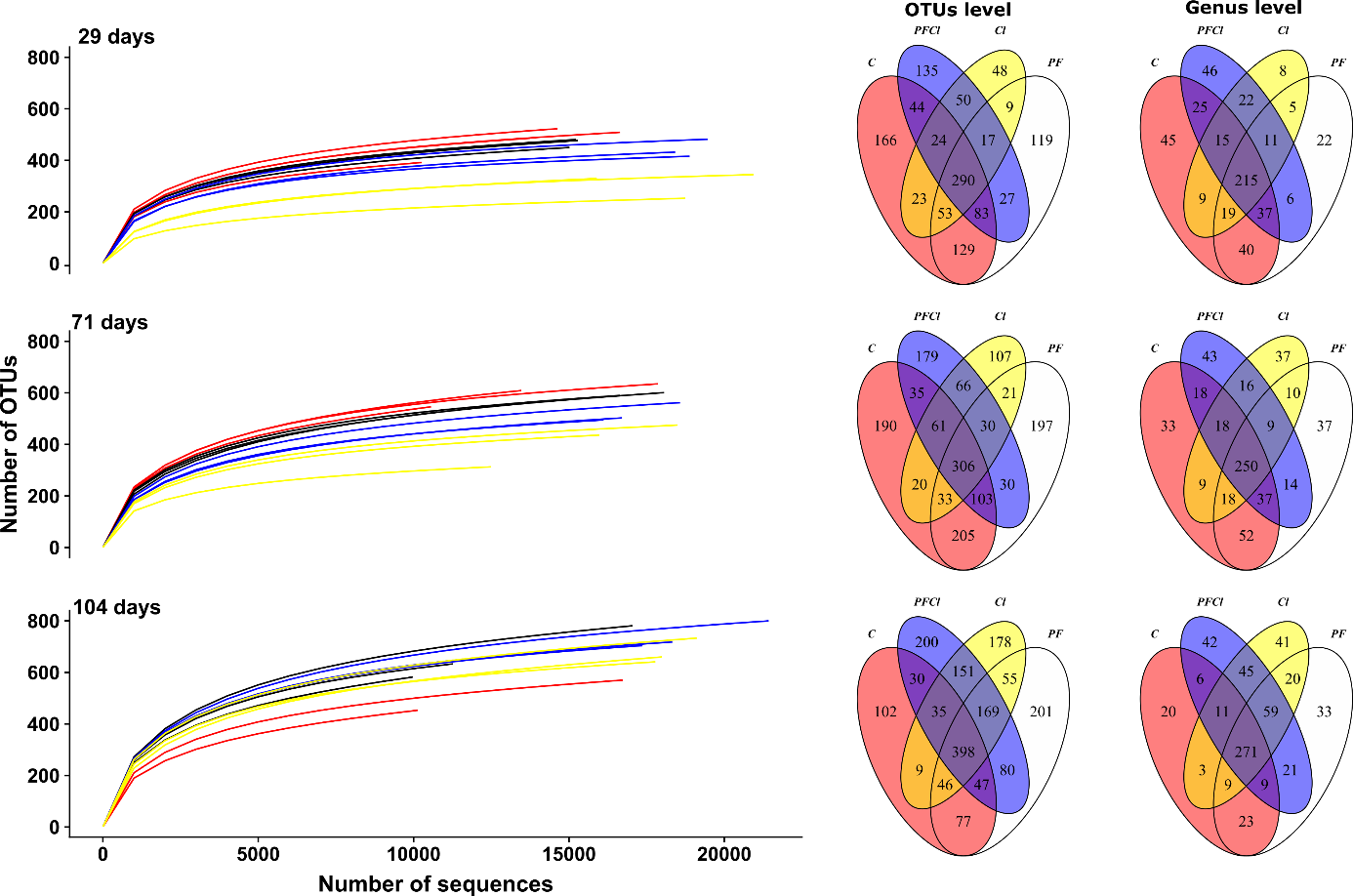

**Figure S2 Rarefaction curves from dripper biofilms and Venn Diagram of bacterial OTUs and bacterial genera in time.** (Control: red, Pressure flushing: white, Pressure flushing & Chlorination: blue, Chlorination: yellow).

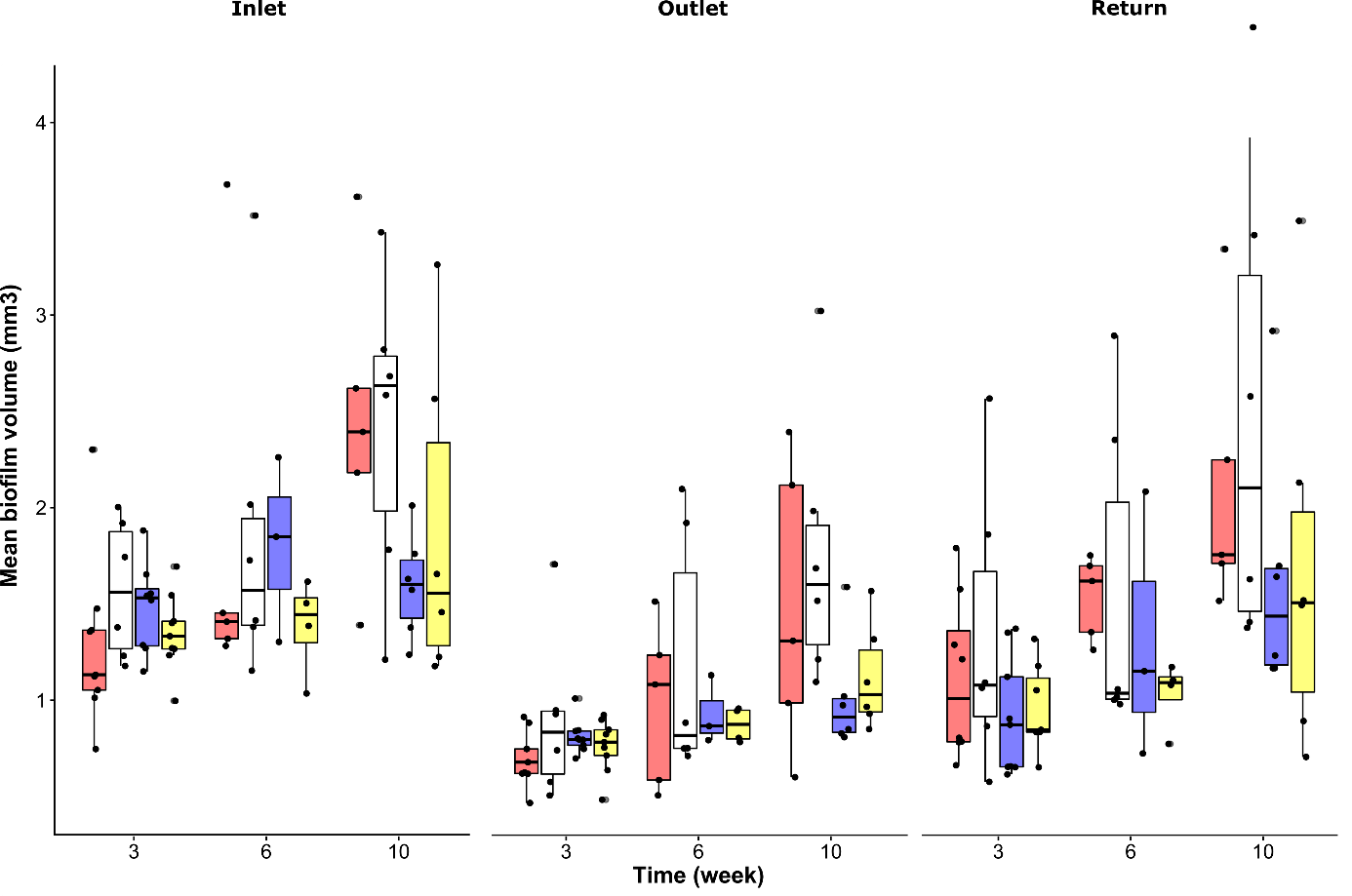

**Figure S3 Biofilm volume during the cleaning period.** Control (●), Pressure flushing (○), Chlorination (●) and Pressure flushing/Chlorination (●).

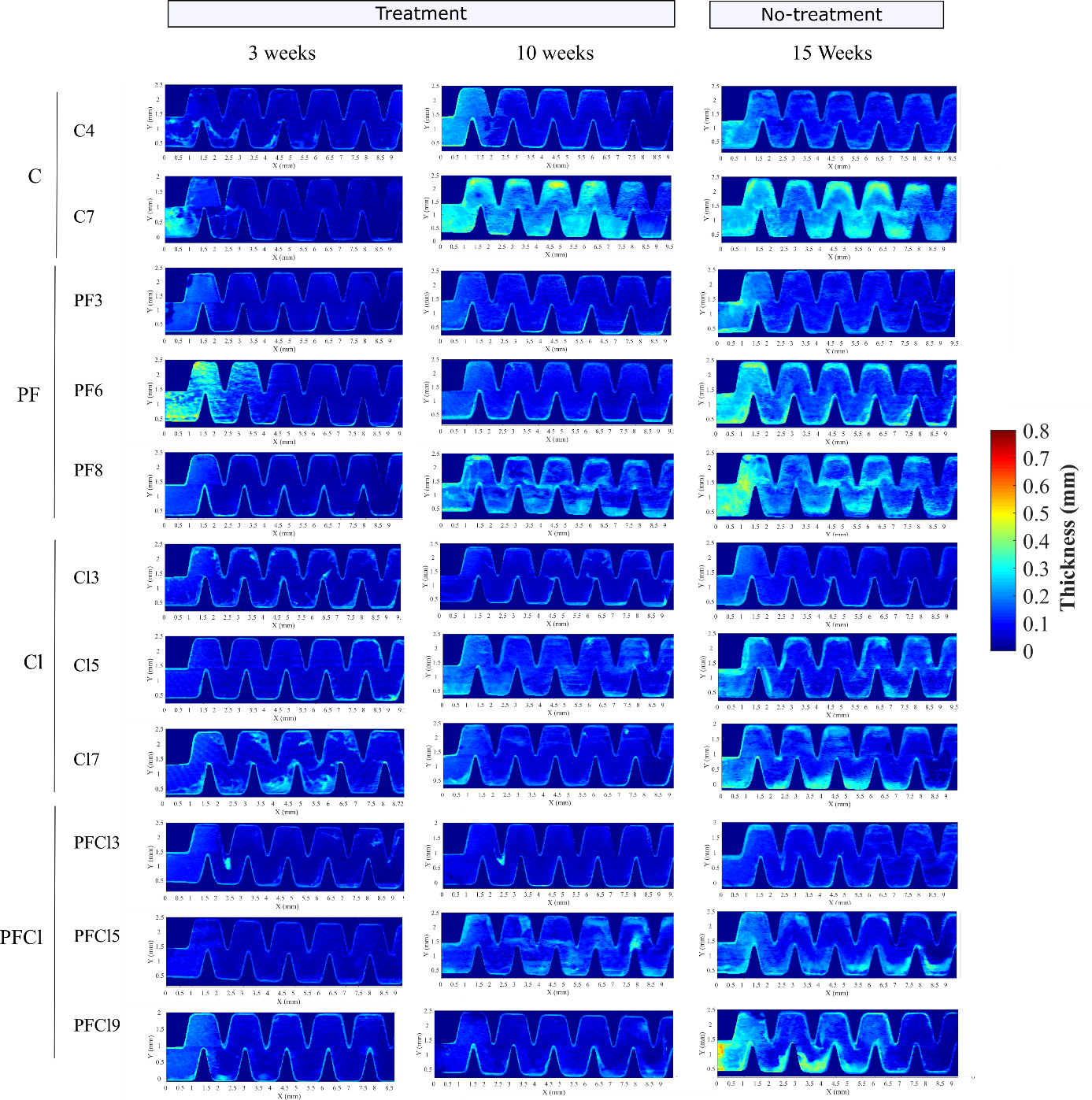

**Figure S4 Biofilm thickness at the outlet of drippers under the C (Control), PF (Pressure flushing), Cl (Chlorination) and PFCl (Pressure flushing/Chlorination) conditions measured at 3, 10 and 15 weeks.** The drippers presented are those also analysed by 16S rRNA sequencing at 104 days.

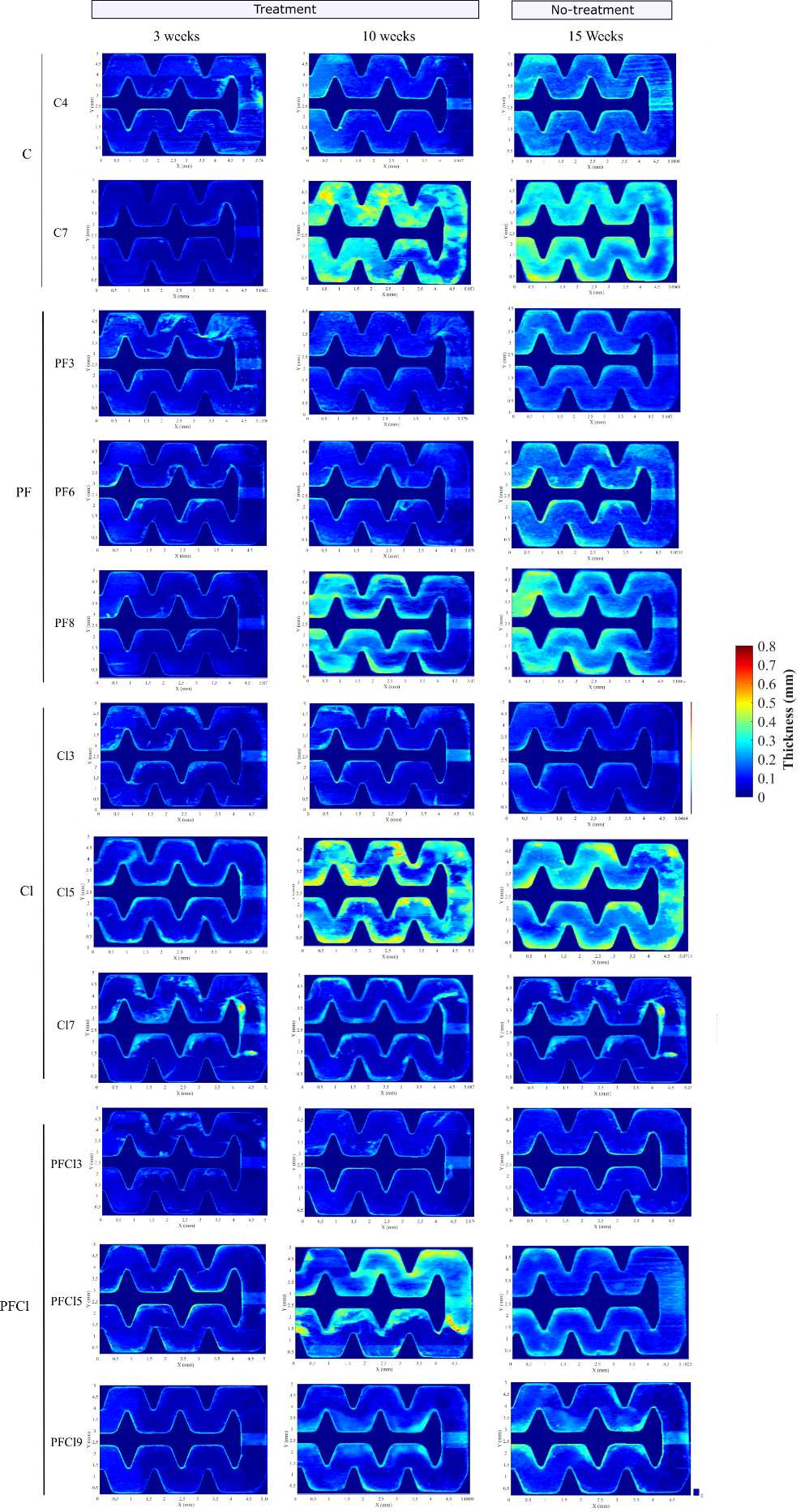

**Figure S5 Biofilm thickness at the return of drippers under the C (Control), PF (Pressure flushing), Cl (Chlorination) and PFCl (Pressure flushing/Chlorination) conditions measured at 3, 10 and 15 weeks.** The drippers presented are those also analysed by 16S rRNA sequencing at 104 days.

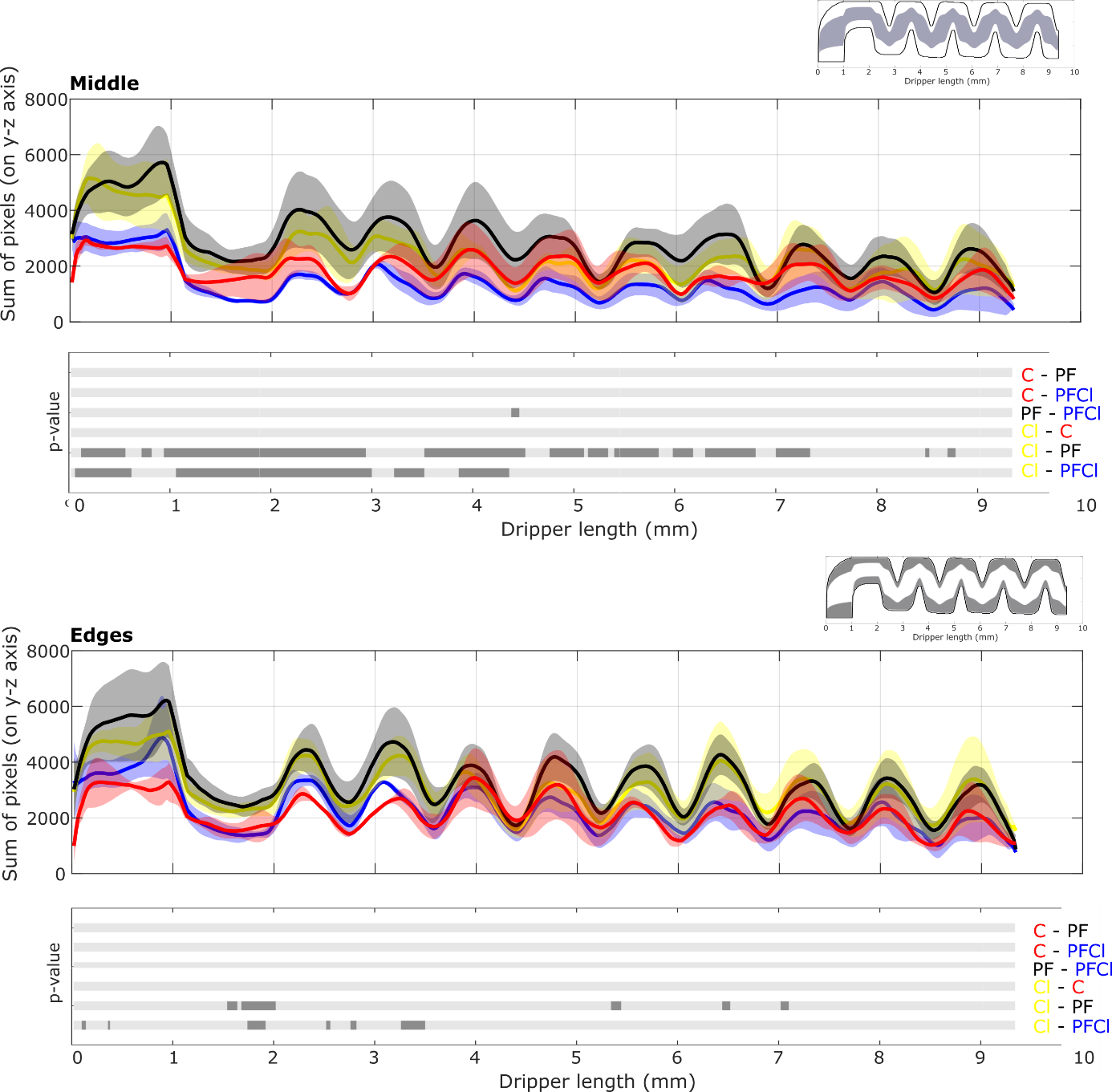

**Figure S6 Mean number of pixels associated with the biofilm mass (and standard deviation) in the middle and edges of the inlet dripper channel after one month without cleaning.** Control (C-●), Pressure flushing (PF-●), Chlorination (Cl-●) and Pressure flushing combined with Chlorination (PFCl-●); n=6 per condition. P-value graphs show the results of the Wilcoxon tests with ■: non-significant, ■: p<0.1, ■: p<0.05.

**Table S1** **Top abundant genera (<3%, x10^-1^) of genera found specifically in one dripper type for each sampling time.–un: unclassified, _ge: genus**

| **Time (days)** | **Control (C)** | | **Pressure flushing (PF)** | | **Chlorination & Pressure flushing (PFCl)** | | **Chlorination (Cl)** | |
| --- | --- | --- | --- | --- | --- | --- | --- | --- |
|  | **Genus** | **Relative abundance** | **Genus** | **Relative abundance** | **Genus** | **Relative abundance** | **Genus** | **Relative abundance** |
| 29d | Comamonadaceae_un | 6.8 ± 1.2 | Comamonadaceae_un | 7.3 ± 2.1 | Comamonadaceae_un | 21.7 ± 7.4 | *Azospira* | 19.2 ± 6.2 |
|  | env.OPS_17_ge | 6.1 ± 2.2 | *Dechloromonas* | 6.7 ± 1.3 | *Flavobacterium* | 9.6 ± 1.3 | *Cloacibacterium* | 14.7 ± 4.1 |
|  | *Fusibacter* | 5.7 ± 1 | env.OPS_17_ge | 5.1 ± 1.3 | *Dechloromonas* | 5.6 ± 3.1 | Comamonadaceae_un | 13.9 ± 3 |
|  | *Azospira* | 5 ± 1.6 | *Terrimonas* | 4.6 ± 1 | env.OPS_17_ge | 5.5 ± 3 | *Flavobacterium* | 12.4 ± 4.7 |
|  | *Pseudomonas* | 4.9 ± 0.4 | *Fusibacter* | 4.4 ± 0.2 | *Cloacibacterium* | 4.9 ± 1.3 | *Pseudoxanthomonas* | 5.1 ± 3 |
|  | *Dechloromonas* | 4.9 ± 1.3 | *Pseudomonas* | 4.4 ± 0.9 | *Pseudomonas* | 4.4 ± 1.6 | *Dechloromonas* | 4.7 ± 2 |
|  | *Terrimonas* | 4.1 ± 1.5 | *Flavobacterium* | 3.7 ± 0.5 |  |  | Comamonadaceae_un | 15.2 ± 1.1 |
| 71d | uncultured_ge | 6.5 ± 1.1 | *Terrimonas* | 7.2 ± 2.2 | Comamonadaceae_un | 22.5 ± 3.8 | *Azospira* | 8 ± 1.1 |
|  | *Terrimonas* | 5.7 ± 0.9 | *Dechloromonas* | 6.5 ± 2.5 | *Fusibacter* | 8.1 ± 3.6 | *Flavobacterium* | 6.3 ± 0.9 |
|  | *Denitratisoma* | 5.4 ± 0.4 | uncultured_ge | 6.3 ± 3 | *Azospira* | 4.8 ± 1.2 | *Fusibacter* | 5.8 ± 1.1 |
|  | *Dechloromonas* | 4.1 ± 0.7 | Comamonadaceae_un | 5.3 ± 3 | *Pseudomonas* | 4.5 ± 0.4 | WCHB1-32_ge | 4.7 ± 0.1 |
|  | Run-SP154_ge | 4.1 ± 1.2 | *Denitratisoma* | 5.2 ± 0.1 |  |  | *Paludibacter* | 4.5 ± 0.8 |
|  | Comamonadaceae_un | 3.6 ± 0.6 | uncultured | 3.9 ± 0.7 |  |  |  |  |
|  |  |  | Run-SP154_ge | 3.6 ± 0.8 |  |  |  |  |
| 104d | *Dechloromonas* | 12.3 ± 3.5 | *Dechloromonas* | 5.8 ± 1.4 | Comamonadaceae_un | 9.1 ± 1 | Comamonadaceae_un | 14.3 ± 2.2 |
|  | *Aquabacterium* | 8.4 ± 0.3 | *Denitratisoma* | 5.1 ± 0.6 | *Dechloromonas* | 7.4 ± 2.2 | Coriobacteriaceae_un | 7.8 ± 0 |
|  | *Herpetosiphon* | 7 ± 0 | *Competibacter* | 4.5 ± 1.2 | uncultured | 6.8 ± 1.4 | uncultured_ge | 4.5 ± 0.8 |
|  | Comamonadaceae_un | 6.6 ± 0.7 | uncultured | 4 ± 0.6 | uncultured_ge | 4.8 ± 0.1 | Christensenellaceae_un | 3.9 ± 0 |
|  | Coriobacteriaceae_un | 6.5 ± 0 | uncultured_ge | 4 ± 0.8 | Christensenellaceae_un | 3.5 ± 0 | *Thiocapsa* | 3.6 ± 0 |
|  | uncultured | 5.8 ± 1 | Run-SP154_ge | 3.9 ± 0.5 |  |  | Run-SP154_ge | 3.3 ± 0.1 |
|  |  |  | Candidatus_Nitrotoga | 3.8 ± 0.4 |  |  | uncultured | 3.2 ± 0.2 |
|  |  |  | *Terrimonas* | 3.8 ± 0.5 |  |  |  |  |

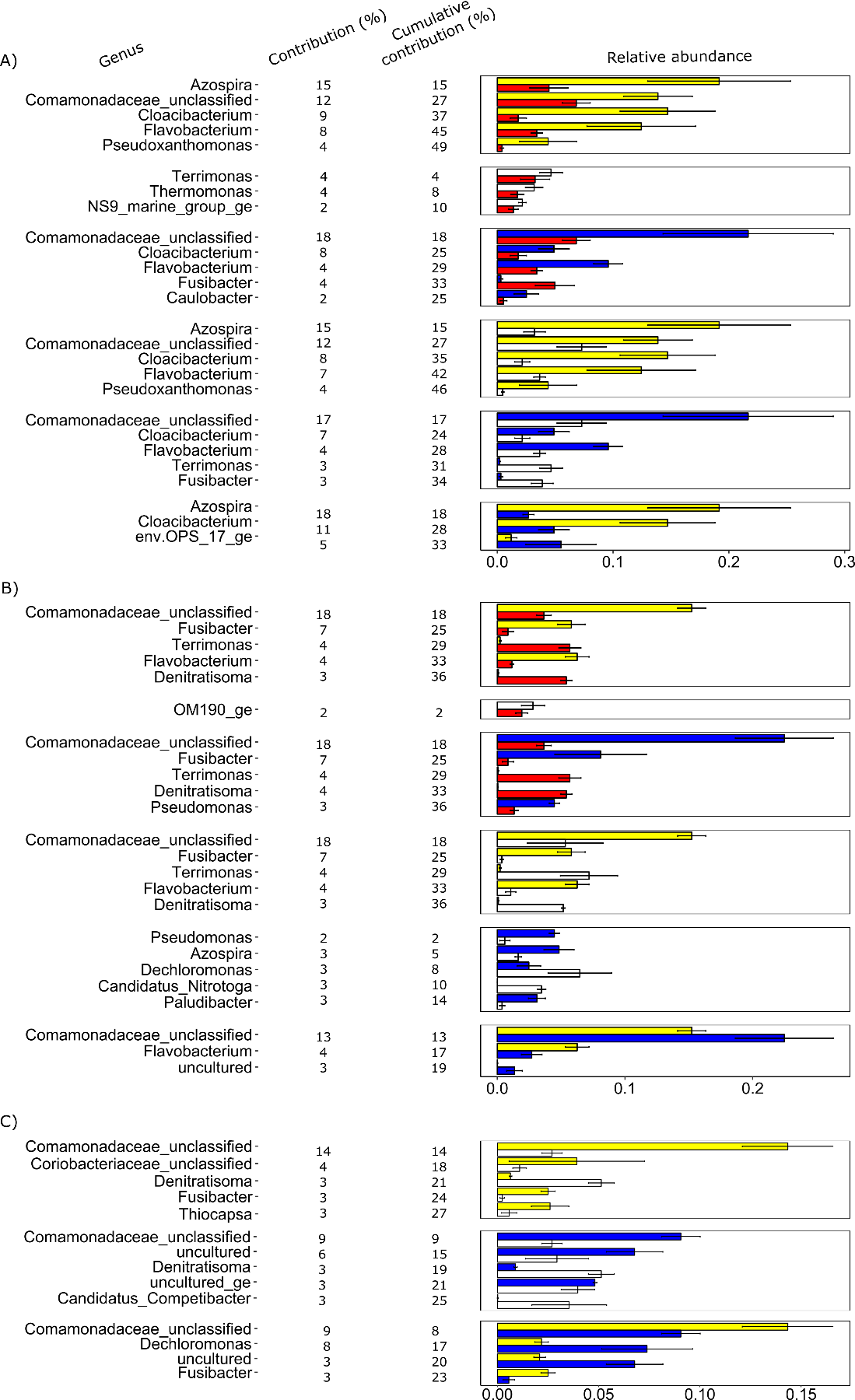

**Figure S8 Most influential bacterial genera in discriminating between 2 conditions (SIMPER analysis).** Control (●), Pressure flushing (○), Chlorination (●) and Pressure flushing/Chlorination (●).

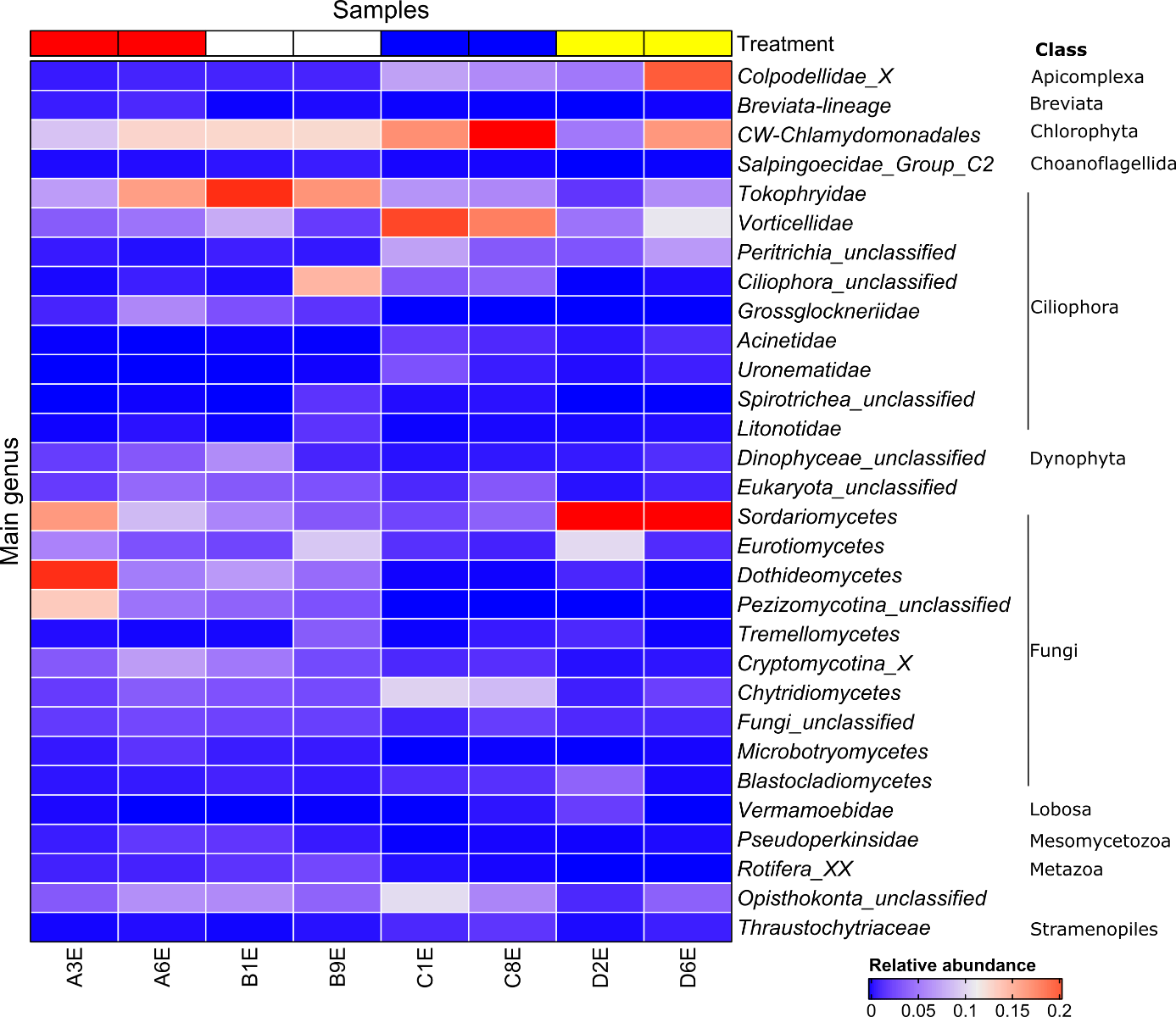

**Figure S9 Heat map of eukaryotes genera from dripper biofilm**s Genera in top thirty relative abundance are shown. Control (●), Pressure flushing (○), Chlorination (●) and Pressure flushing/Chlorination (●).
